# *In-utero* rescue of neurological dysfunction in a mouse model of Wiedemann-Steiner syndrome

**DOI:** 10.1101/2024.07.19.604339

**Authors:** Tinna Reynisdottir, Kimberley Jade Anderson, Andrew Brinn, Katie Franklin, Juan Ouyang, Asbjorg Osk Snorradottir, Cathleen M. Lutz, Aamir R. Zuberi, Valerie Burke DeLeon, Hans Tomas Bjornsson

**Affiliations:** Louma G. Laboratory of Epigenetic Research, Faculty of Medicine, University of Iceland; Department of Genetics and Molecular Medicine, Landspitali University Hospital, Reykjavik, Iceland; Department of Anthropology, University of Florida; Faculty of Medicine, University of Iceland, Reykjavik, Iceland; Department of Pathology, Landspitali University Hospital, Reykjavik, Iceland; The Jackson Laboratory, Bar Harbor; McKusick-Nathans Departments of Genetic Medicine and Pediatrics, Johns Hopkins University

**Keywords:** KMT2A, genetic rescue, Nestin-Cre, WDSTS, Intellectual disability

## Abstract

Wiedemann-Steiner syndrome (WDSTS) is a rare genetic cause of intellectual disability primarily caused by heterozygous loss of function variants in the gene encoding the histone methyltransferase KMT2A. Prior studies have shown successful postnatal amelioration of disease phenotypes for Rett, Rubinstein-Taybi and Kabuki syndromes, related Mendelian disorders of the epigenetic machinery. To explore whether the neurological phenotype in WDSTS is treatable *in-utero*, we created a novel mouse model carrying a loss of function variant in between two loxP sites. *Kmt2a^+/LSL^* mice demonstrate core features of WDSTS including growth retardation, craniofacial abnormalities, and hypertrichosis as well as hippocampal memory defects. The neurological phenotypes show rescue upon restoration of *KMT2A in-utero* following breeding to a nestin-Cre. Together, our data provide a novel mouse model to explore the therapeutic window in WDSTS. Our work suggests that WDSTS has a window of opportunity extending at least until the mid-point of *in-utero* development, making WDSTS an ideal candidate for future therapeutic strategies.

Graphical abstract

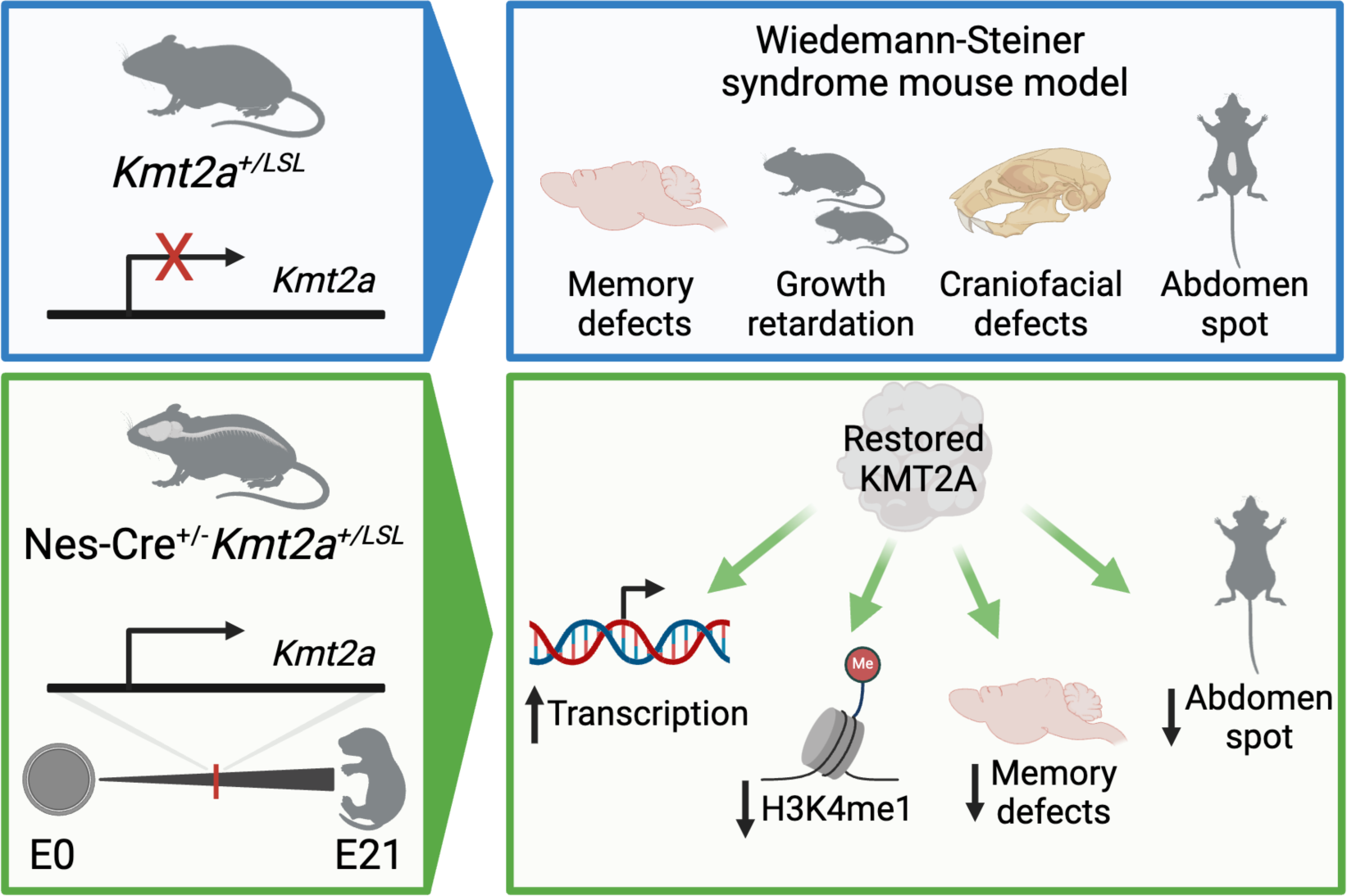

## Introduction

Wiedemann Steiner syndrome (WDSTS, OMIM: #605130) is a Mendelian disorder of the epigenetic machinery (MDEM), inherited in an autosomal dominant manner. WDSTS is primarily caused by *de novo* heterozygous variants in the gene encoding the histone-lysine methyltransferase 2A (KMT2A, previously MLL, MLL-1 or ALL in literature) [1]. KMT2A plays a crucial role in the mono-,di-and trimethylation of lysine 4 of histone 3 (H3K4me1/2/3) [2], modifications generally associated with active gene expression [3], [4], occurring at enhancers and promoters. KMT2A has been shown to be essential for postnatal neurogenesis in mice, and as a regulator of neural progenitor proliferation and differentiation [5], [6]. Key features of WDSTS include intellectual disability, growth retardation, hypotonia, craniofacial defects, gastrointestinal problems, and hypertrichosis. Previous mouse studies have shown postnatal rescue of neurological defects in several other MDEMs, including Rubinstein-Taybi [7], Rett [8] and Kabuki syndromes [9], [10]. To extend these findings to WDSTS, we have created a novel mouse model for WDSTS (*Kmt2a^+/LSL^*). Earlier mouse models have demonstrated homozygous knock-out (KO) of *Kmt2a* (*Kmt2a^-/-^* mice) is lethal, while heterozygous mice *(Kmt2a*^+/-^) exhibited growth retardation, skeletal defects, hemopoietic abnormalities [11] as well as neurological defects in visuospatial-and associative memory [12] and increased aggressive behavior [13]. Notably, it has been shown that joint constitutive KO of the writer-eraser duo KMT2A and KDM5C rescues disease phenotypes seen in either disease model, including rescue of transcriptome abnormalities, neuronal structure and aggressive behavior. This finding suggests that the enzymatic function of KMT2A plays a role in disease and that a specific balance is required among these histone machinery components [13], [14]. In contrast, homozygous mice with a mutation specifically altering the C-terminal enzymatic SET domain remain viable and fertile [15]. In concordance with the latter, our previous work has shown that missense variants causing WDSTS are mainly clustered outside of the enzymatic SET domain, and within the DNA binding CXXC domain [16], suggesting that non-enzymatic roles of KMT2A may be key in WDSTS pathogenesis.

To explore whether WDSTS is treatable *in-utero*, we created and characterized a novel mouse model (*Kmt2a^+/LSL^*) which carries a heterozygous insertion of a loxP-stop-loxP (LSL) cassette in intron 1 of *Kmt2a* resulting in heterozygous KO of the affected allele. The model (*Kmt2a^+/LSL^*) serves as a novel disease model for WDSTS as well as allowing temporal control of *Kmt2a*. By crossing the model to a nestin-Cre model, we demonstrated that *in-utero* rescue of KMT2A levels could rescue WDSTS neurological phenotypes in primary neuronal progenitor cells (NPCs) from these mice and validated this finding by *in vivo* studies.

## Results

### Targeted knock-out of *Kmt2a* recapitulates key features of WDSTS

WDSTS broadly affects patients with common clinical features within and outside the nervous system. To get an unbiased view of WDSTS phenotypes we have summarized available data from an ongoing RARE-X WDSTS collection. The majority of the patients exhibit neurological issues (93.8%), most commonly intellectual disability and/or developmental delay, with a large part exhibiting behavioral and/or psychiatric problems (75.8%). Other common features include growth retardation (95.4%), dental and/or oral problems (79.4%) and hypotonia (60.0%). Distinctive features of WDSTS also include digestive system issues (76.9, most commonly constipation) and hypertrichosis cubiti (57%) (Table 1).

**Table 1.** Defining clinical features of WDSTS patients including percentage and count of patients demonstrating the observed phenotype. Data gathered from RARE-X patient database and literature [1] [17]

| Clinical feature | WDSTS patients % (n) |
| --- | --- |
| Growth retardation | 95.4% (62/65) |
| Brain and/or neurological issues | 93.8% (60/64) |
| Eye and/or vision issues | 84.7% (50/59) |
| Sleep issues | 80.3% (53/66) |
| Dental and/or oral issues | 79.4% (50/63) |
| Digestive system issues | 76.9% (50/65) |
| Behavior and/or psychiatric issues | 75.8% (47/62) |
| Hypotonia | 60.0% (33/55) |
| Hypertrichosis cubiti | 57.0% (57/100) |
| Head, face and/or neck issues | 56.3% (36/62) |

To elucidate the *in-utero* neurological malleability of WDSTS, we created a novel mouse model (*Kmt2a^+/LSL^)* with a heterozygous loxP-stop-loxP (LSL) cassette insertion in intron 1 of *Kmt2a* (Figure 1A). The LSL cassette induces an early poly-A signal leading to immediate truncation of the mRNA transcript in intron 1 of the affected allele, allowing rescue of expression with exposure to Cre-recombinase. The mouse model was created via a service contract with Jackson Laboratory who tested functionality of the cassette mice by breeding *Kmt2a^+/LSL^* mice to a Sox2-Cre containing mouse model which demonstrated that loxP sites were functional (see Methods). To further verify the orientation and completeness of the cassette, we performed Oxford nanopore technology (ONT) long-read sequencing on genomic DNA (gDNA) from *Kmt2a^+/LSL^* mice, from which single reads revealed the entirety of the cassette integrated into intron 1 of *Kmt2a* in the correct orientation (Figure 1A). Since this experiment provides whole-genome coverage we also interrogated two predicted potential off-target sites from cassette integration and observe no obvious variants or indels at these sites (Supplemental Figure 1A-D). The offspring of *Kmt2a^+/LSL^* mice show skewing of the expected Mendelian ratio with a significant bias against the inheritance of the mutant allele (p < 0.01, chi-squared test), with 40.9% (90/220) *Kmt2a^+/LSL^* mice born and 59.1% (130/220) *Kmt2a^+/+^* wild-type littermates. Similar to other *Kmt2a*-deficient models [11], *Kmt2a^+/LSL^* mice demonstrate lethality in homozygosity, with no *Kmt2a^LSL/LSL^* mice born (Supplemental Figure 1E-F). As predicted, the *Kmt2a^+/LSL^* mice demonstrate ∼50% reduction of *Kmt2a* mRNA transcripts as quantified by real-time quantitative polymerase chain reaction (RT-qPCR) (n=3-7, Figure 1B). In addition to observing appropriate integration of the LSL cassette, we observed characteristic WDSTS phenotypes of the *Kmt2a^+/LSL^* mice, supporting the notion that these mice adequately recapitulate the human disease. As previously reported in *Kmt2a*^+/-^ mice [11], *Kmt2a^+/LSL^* mice exhibit growth deficiency with reduced weight compared to *Kmt2a^+/+^* littermates (Figure 1C, Supplemental Figure 2A-B), alongside hemopoietic abnormalities similar to what has been previously described [11], including a decreased lymphocyte percentage, increase in monocyte percentages and an increase in platelet distribution width (Supplemental Figure 3). Given the consistent description of hypertrichosis in individuals with WDSTS [17], we decided to explore the skin histology of newborn P1 (postnatal day 1) pups. *Kmt2a^+/LSL^* mice show increased hair follicle count on the abdomen compared to their *Kmt2a^+/+^* littermates, while there was no significant increase in hair follicle count on their back (Figure 1D). Furthermore, we did not observe any obvious differences in hair regrowth upon removal of hair from the backs of the *Kmt2a^+/LSL^* mice compared to their wild-type littermates (Supplemental Figure 2C). This may indicate that the basis of hypertrichosis in WDSTS may involve increased number of hair follicle in contrast to increased rate of hair regrowth. Since severe constipation and other GI symptoms are a common phenotype in WDSTS patients, we stained the distal colon sections of *Kmt2a^+/+^* and *Kmt2a^+/LSL^* mice for Calretinin, a common diagnostic marker for Hirschsprung’s disease (HD) that marks ganglion cells [18]. Our results showed positive staining in both genotypes (Supplemental Figure 2D), indicative of normal presence of ganglions and nerve fibers in our WDSTS model. However, this does not rule out more subtle abnormalities of the enteric nervous system or problems relating to smooth muscle of the intestines.

**Figure 1:**
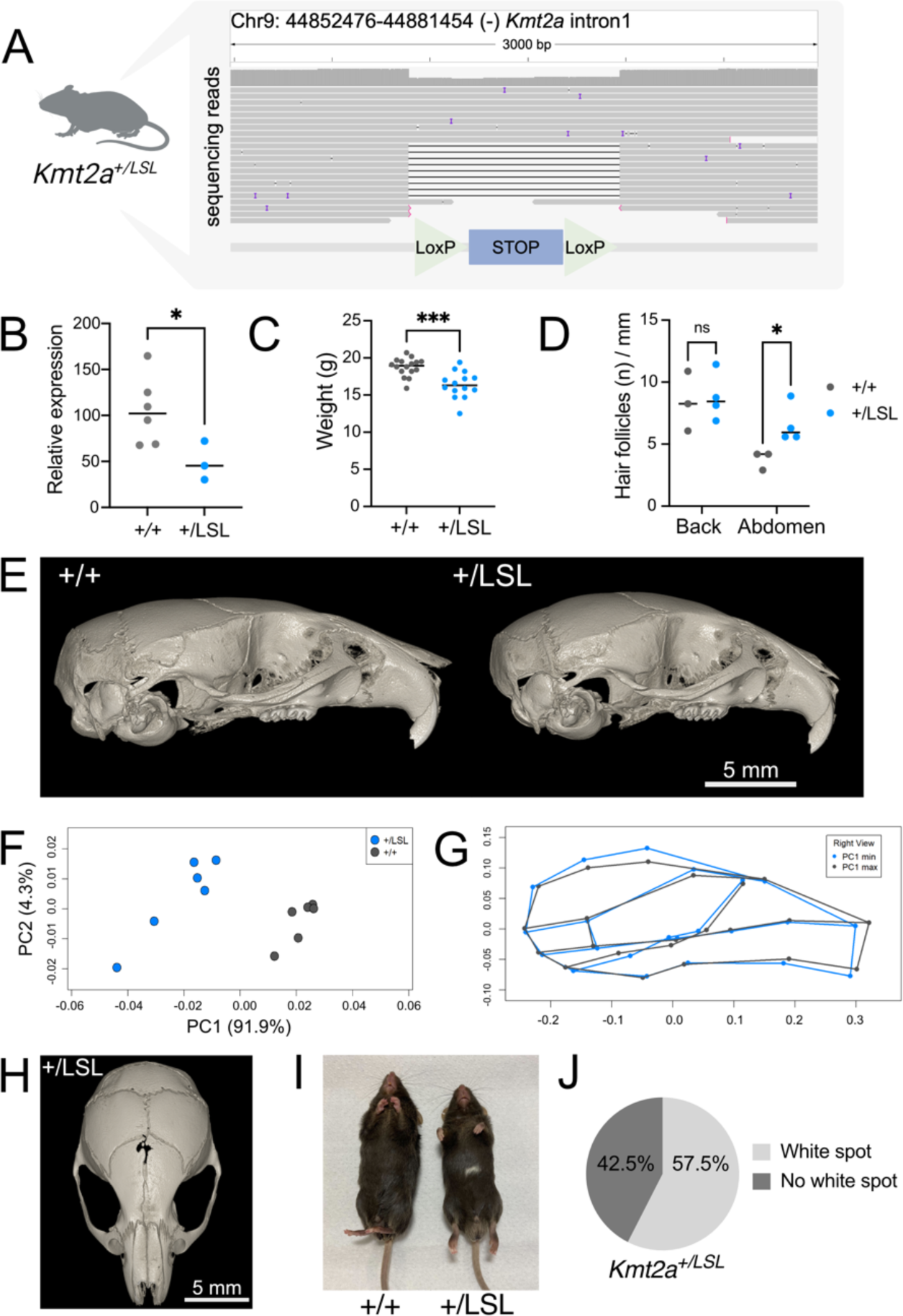
A novel Wiedemann-Steiner syndrome mouse model (*Kmt2a^+/LSL^*) recapitulates core phenotypes seen in individuals with WDSTS. (A) Schematic figure of *Kmt2a^+/LSL^* mouse model with a heterozygous loxP-stop-loxP cassette inserted in intron 1 of *Kmt2a* showing ONT sequencing reads of the cassette. Each line represents an individual long read including several that span the entire cassette as well as boundaries on either side allowing verification of its orientation. (B) Relative total *Kmt2a* mRNA expression is decreased in *Kmt2a^+/LSL^* mice compared to *Kmt2a^+/+^*littermates (n=7 *Kmt2a^+/+^* and n= 3 *Kmt2a^+/LSL^*). (C) *Kmt2a^+/LSL^* mice weigh significantly less compared to *Kmt2a^+/+^* littermates (n=14-16 per genotype, 8-week females). (D) Hair follicle count per mm of skin is decreased on abdomen but not on the back of newborn *Kmt2a^+/LSL^* pups compared to littermates (n=3-4 per genotype). (E) Representative CT images of *Kmt2a^+/+^*and *Kmt2a^+/LSL^* mice from lateral view, the depicting the craniofacial phenotype. (F) PCA plot of the craniofacial structure showing a distinct separation of the genotypes along PC1. (G) The *Kmt2a^+/LSL^*mice are positioned at the negative end of PC1 and characterized by increased height and width of the neurocranium, a shortened midface, and ventral bowing of the cranium. (H) Representative figure of the gap of the interfrontal suture present in the *Kmt2a^+/LSL^* mice. (I) Representative figure of the white abdominal spot of a *Kmt2a^+/LSL^*mouse next to a wild-type littermate. (J) Majority of *Kmt2a^+/LSL^*mice (57.5%, n=40) present with a white abdominal spot, a phenotype not present in *Kmt2a^+/+^* littermates. ns: not significant, *p< 0.05, **p < 0.01, ***p < 0.001, ****p < 0.0001.

To evaluate whether craniofacial abnormalities are present, we performed CT scans on the *Kmt2a^+/LSL^* mice (n=6) mice and their *Kmt2a^+/+^* littermates (n=6). A Procrustes ANOVA yielded a significant effect of genotype on shape (Figure 1E, p < 0.001). Principal Components Analysis (PCA) showed a clear separation between the groups along PC1, which describes 91.9% of the symmetric shape variance (Figure 1F). The *Kmt2a^+/LSL^* mice are characterized by having increased height and width of the neurocranium and ventral bowing of the cranium compared to littermates (Figure 1G). Additionally, we observed a prominent gap at the interfrontal suture in the *Kmt2a^+/LSL^* mice, which was not observed in any of the *Kmt2a^+/+^* littermates (Figure 1H). Similar craniofacial defects observed in Kabuki syndrome have been linked to neural crest defects [19], [20], [21]. Given the shared characteristics of these syndromes, it raises the question whether neural crest defects could also contribute to abnormalities found in WDSTS. Interestingly, in our model, a majority (57.5%) of the *Kmt2a^+/LSL^* mice present with a white abdominal spot, ranging from few distinct hairs to spots ∼1-2 cm in length, most common along the midline (Figure 1I-J, Supplemental Figure 2E), suggestive of defects in the melanoblast migration from the neural crest [22]. In contrast, we have never observed such a white spot in wild-type littermates.

### Observed neurological defects in the *Kmt2a^+/LSL^* mouse model

Consistent with the hypotonia observed in some WDSTS patients, *Kmt2a^+/LSL^* mice exhibit significantly longer (p-value < 0.01, unpaired t-test) surface righting time compared to *Kmt2a^+/+^* littermates, indicating of lack of core strength and coordination (Figure 2A) and a significant decrease (p-value < 0.01, unpaired t-test) in hindlimb suspension time compared to *Kmt2a^+/+^*, indicative of increased fatigue in hindlimbs (Figure 2B), tested on postnatal day 6 (PD6, n=14-27). The visuospatial memory function in adult *Kmt2a^+/LSL^* mice was tested in an open Y maze (Figure 2C). The *Kmt2a^+/LSL^* mice did not exhibit a significant decrease of arms entered during the test time (p-value = 0.1576, unpaired t-test), suggesting the adult mice do not exhibit a gross motor phenotype (Figure 2D). However, the *Kmt2a^+/LSL^* mice demonstrate significant deficits (p-value < 0.01, unpaired t-test) in spontaneous alternations compared to *Kmt2a^+/+^* littermates, indicative of defective working memory reliant on the hippocampus and prefrontal cortex (n=18-24 per genotype, Figure 2E) [23], [24]. These findings are in concordance of previous studies describing the importance of *Kmt2a* for memory consolidation [12]. Histologically, when we measured the size of the granule cell layer of the dentate gyrus (DG) of the hippocampus with DAPI staining, the total area was significantly smaller (p-value < 0.05, unpaired t-test) in the *Kmt2a^+/LSL^* mice, compared to *Kmt2a^+/+^* littermates, when corrected for body weight (n=3-4 per genotype, Figure 2F-H). A decreased size of the dentate gyrus has been previously described in a mouse model of Kabuki syndrome [9] and later validated in individuals with Kabuki syndrome [25]. Taken together, our findings support that the *Kmt2a^+/LSL^* mouse model recapitulates core human WDSTS disease phenotypes and allows for quantitative measurement of a number of metrics related to the neurological phenotype that can be used to investigate the *in-utero* malleability.

**Figure 2:**
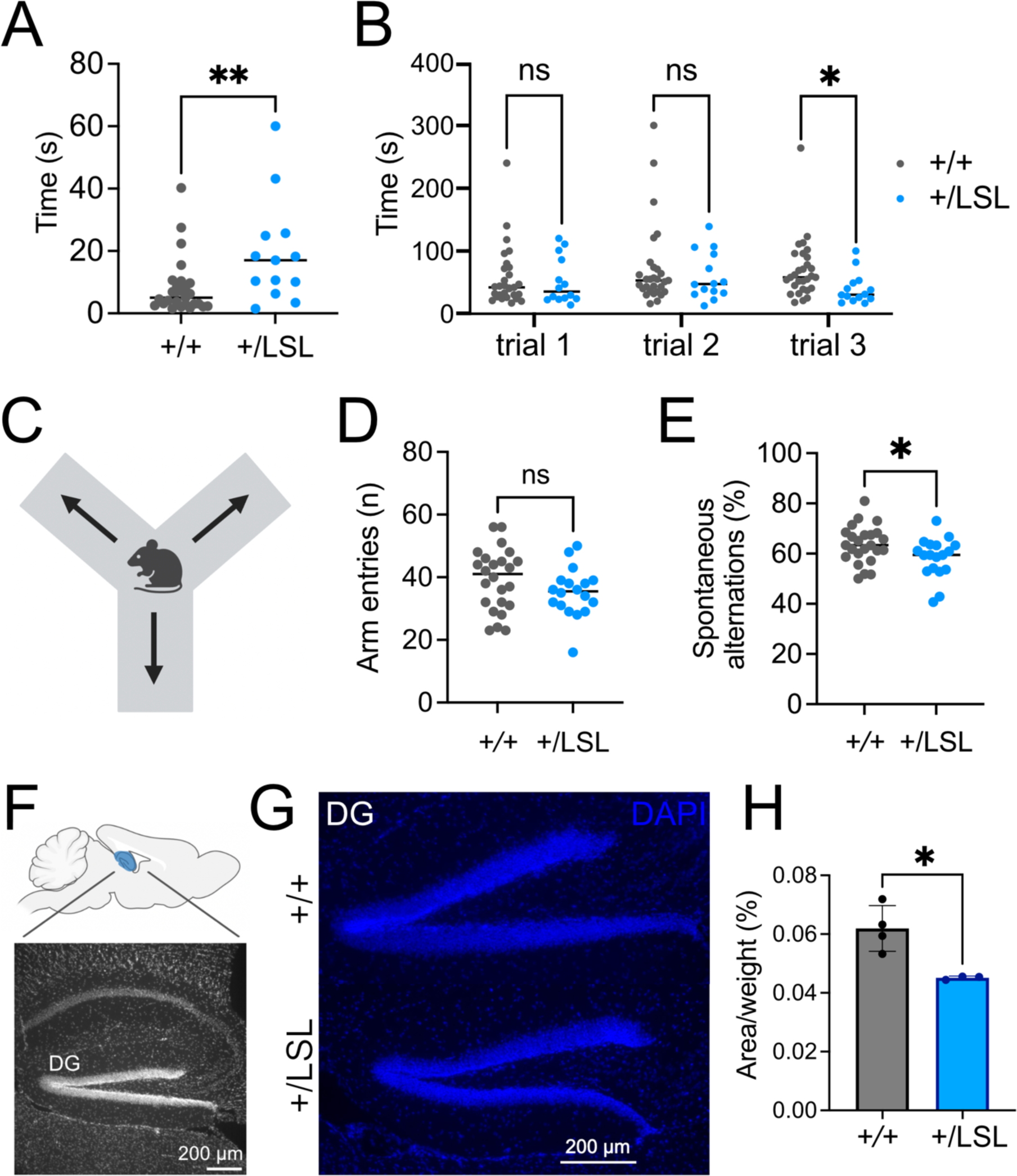
Neurological defects observed in the *Kmt2a^+/LSL^* mouse model. (A) Increased surface righting time in *Kmt2a^+/LSL^*and (B) decreased hindlimb suspension time at the third trial in hypotonia tests (PD6, n= 27-29, *Kmt2a^+/+^*, n=14-15, *Kmt2a^+/LSL^*). (C) Schematic overview of a Y maze. A spontaneous alternation is completed when mouse has entered each arm of the maze consecutively. (D) Adult *Kmt2a^+/LSL^* mice do not show a significant decrease of total arm entries in a Y maze (n= 24, *Kmt2a^+/+^*, n= 18, *Kmt2a^+/LSL^*). (E) Adult *Kmt2a^+/LSL^* mice exhibit a significant decrease in percentage of spontaneous alternations in a Y maze. (F) Schematic figure of the mouse hippocampus with a representative image showing the DG region of the hippocampus. (G) Representative images of a DAPI staining of the DG area of hippocampus in *Kmt2a^+/+^* and *Kmt2a^+/LSL^* mice. (H) Quantification of granule cell layer area of the DG demonstrating a decrease in *Kmt2a^+/LSL^* mice compared to wild-type littermates when corrected for body weight (n=3-4 per genotype). ns: not significant, *p< 0.05, **p < 0.01, ***p < 0.001, ****p < 0.0001.

### Nes-Cre^+/-^*Kmt2a^+/LSL^* mice exhibit successful cassette removal

To test *in-utero* rescue of the neurological phenotypes, we crossed our *Kmt2a^+/LSL^* model with a nestin-Cre model (Nes-Cre^+/+^) which should allow *in-utero* rescue of *Kmt2a* expression in a cell-type specific manner in the nervous system at mid to late gestation (Figure 3A). We hypothesize that with a knock in of *Kmt2a* expression in nervous tissue, the markers of neurological dysfunction will show rescue in Nes-Cre^+/-^*Kmt2a^+/LSL^* mice, with a prior study [13] demonstrating rescue at the time of conception. To validate the efficacy of the LSL cassette recombination in Nes-Cre^+/-^*Kmt2a^+/LSL^* mice in the nervous system we performed PCR with primers targeting the LSL cassette in the hippocampus and skin tissue from the ear using gDNA and compared to Nes-Cre^+/-^*Kmt2a^+/+^* controls. We observe an obvious decrease of the amount of LSL cassette in the nervous tissue but not in the skin tissue, of Nes-Cre^+/-^*Kmt2a^+/LSL^* mice compared to controls (Supplemental Figure 4). To further validate the removal of the LSL cassette and the ability to regain normal *Kmt2a* expression, we quantified the relative mRNA expression in whole brain lysates of P1 pups and observed no significant difference between Nes-Cre^+/-^*Kmt2a^+/+^* and Nes-Cre^+/-^*Kmt2a^+/LSL^* mice (Figure 3B, n=3 per genotype) in contrast to a significant difference between the genotypes without the Cre (Figure 1B).

**Figure 3:**
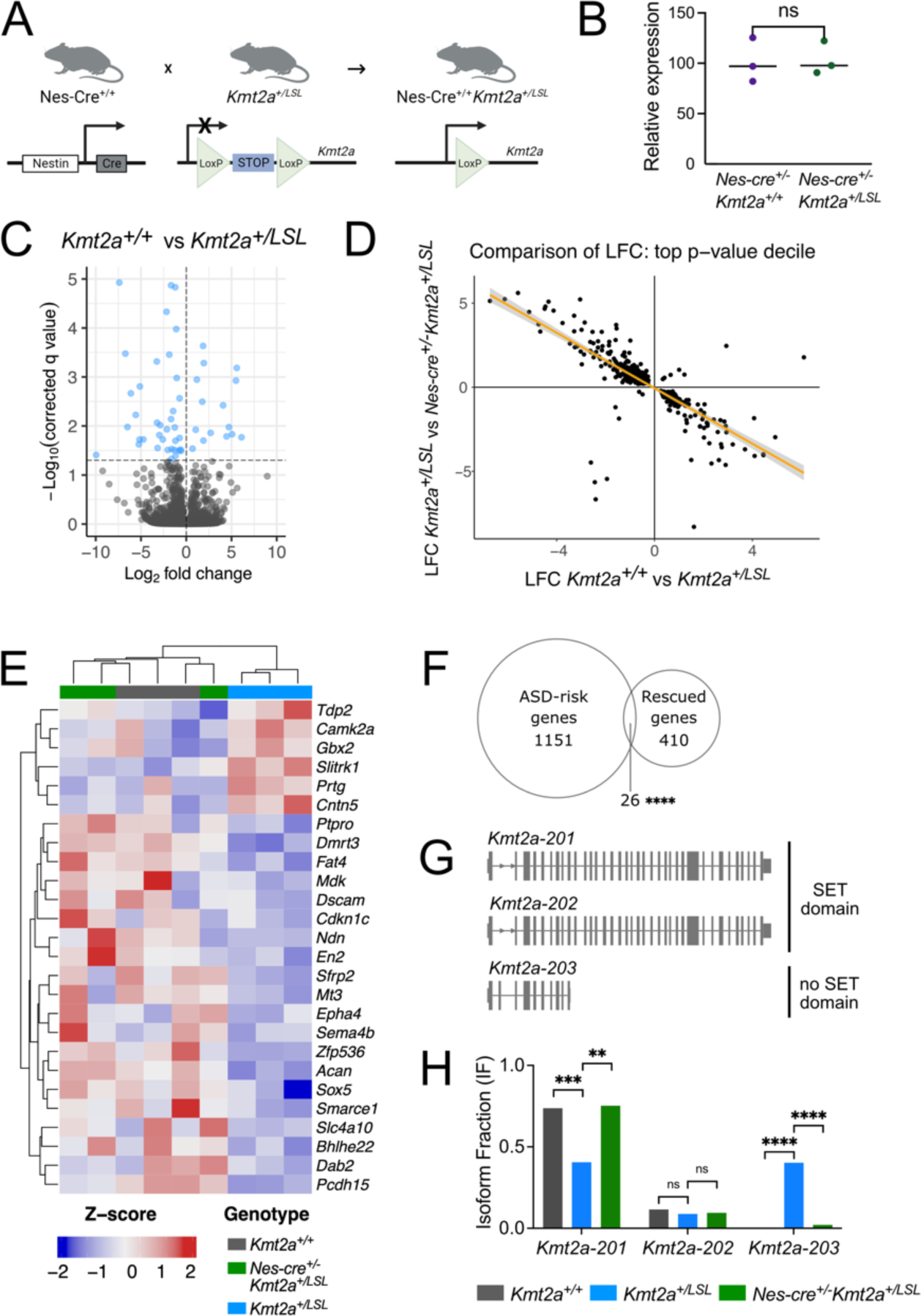
Transcriptional consequences of *Kmt2a* loss and rescue. (A) A schematic overview of the nestin-Cre strategy. By exposing the *Kmt2a^+/LSL^* mice to nestin-Cre *Kmt2a* levels are expected to be restored *in-utero* in the nervous system of the Nes-Cre^+/-^*Kmt2a^+/LSL^*mice *in-utero* allowing us to test *in-utero* rescue. (B) RT-qPCR demonstrates that relative *Kmt2a* mRNA expression is unchanged in Nes-Cre^+/-^*Kmt2a^+/LSL^* mice compared to Nes-Cre^+/+^*Kmt2a^+/+^* littermates (n=3 per genotype) in nervous tissue. (C) Volcano plot showing the log2 fold changes of differentially expressed genes comparing *Kmt2a^+/+^* vs *Kmt2a^+/LSL^* mNPCs with significant genes labelled in blue. (D) Correlation plot showing the negative correlation between log2 fold changes between *Kmt2a^+/+^* vs *Kmt2a^+/LSL^* and then between *Kmt2a^+/LSL^* vs Nes-Cre^+/-^*Kmt2a^+/LSL^* mice (R^2^=0.57988). (E) Heatmap showing clustering of *Kmt2a^+/LSL^* samples compared to *Kmt2a^+/+^*and Nes-Cre^+/-^*Kmt2a^+/LSL^* samples of the 26 rescued genes from significantly overrepresented biological pathways (with FDR<0.05). Expression values were normalized to z-scores and plotted, with increased expression represented in red and decreased expression in blue. (F) Venn diagram of overlapping rescued genes with 1177 known ASD-risk factor genes, with 26 genes overlapping. (G) Representative figure of *Kmt2a* isoforms and their exons. (H) Isoform switch analysis reveals a shift in isoform fraction (IF) in *Kmt2a^+/LSL^* mNPCs from the longer SET domain-containing *Kmt2a* transcript to the short non-SET domain containing *Kmt2a* transcript. ns: not significant, *p< 0.05, **p < 0.01, ***p < 0.001, ****p < 0.0001.

### Gene expression abnormalities are rescued in Nes-Cre^+/-^*Kmt2a^+/LSL^* mNPCs

To evaluate rescue of transcriptional abnormalities caused by heterozygous *Kmt2a*-deficiency in a homogenous cell population, we dissected the hippocampus from newborn pups (P0) and cultured mouse neuronal progenitor cells (mNPCs). We performed RNA-sequencing using mRNA from mNPCs from both genotypes to identify gene expression changes between *Kmt2a^+/+^* and *Kmt2a^+/LSL^* NPCs (n=3 per genotype). We observed 64 significant differentially expressed genes (DEGs) when comparing *Kmt2a^+/+^* and *Kmt2a^+/LSL^* samples with a majority of genes downregulated (47/64 genes), in concordance with the loss of a transcriptional activator (Figure 3C, Supplemental table 1).

Next, we wanted to assess whether there was an *in-utero* reversal of the abnormalities we uncovered in the RNA-seq dataset. First, we observe a significant overlap of 438 genes (p-value < 0.0001, Fisher’s exact test) within the top-decile of p-values for comparisons between both *Kmt2a^+/+^* vs *Kmt2a^+/LSL^* and *Kmt2a^+/LSL^* vs Nes-Cre^+/-^*Kmt2a^+/LSL^*. Secondly, there is a striking negative correlation of the log_2_-fold change (LFC) of these genes between the comparisons, where 96.9% of genes (423/438 genes) demonstrated such an effect (R^2^=0.57988, Figure 3D, Supplemental table 2). In essence, these results indicate that downregulated genes from *Kmt2a^+/LSL^* vs *Kmt2a^+/+^* show upregulation in *Kmt2a^+/LSL^* mice in presence of the nestin-Cre and vice versa. We performed an over-representation analysis (ORA) of the genes showing such rescue and observed significant overrepresentation of pathways implicated in neuron development and neuron differentiation (Supplemental Figure 5A). We pooled the genes from the overrepresented pathways together and can observe a similar trend, with 20/26 genes downregulated in *Kmt2a^+/LSL^* samples (Figure 3E, see filtering criteria in Methods).

Notably, we observed down-regulation of a transcription factors, such as *En2* and *Sox5,* in *Kmt2a^+/LSL^* mNPCs compared to the *Kmt2a^+/+^* mNPCs. Disruption of these transcription factors has previously been linked to defects in hippocampal neurogenesis [26], [27] as well as being associated with intellectual disability and autism-spectrum-disorder (ASD) [28][29]. Since up to 50% of WDSTS patients have ASD [30], we compared the rescued genes to known ASD-risk genes [31] and discovered a significant overlap of 26 genes (p-value < 0.0001, Fisher’s exact test, Figure 3F). Interestingly, *Kmt2a* was not among the significant DEGs in this dataset. However, upon further examination of individual exons in *Kmt2a^+/LSL^* cells, the majority of reads were limited to the first exon of *Kmt2a*, with an overall decrease in coverage for the full length *Kmt2a* transcript (Supplementary figure 5B-C). We then preformed an isoform-specific analysis that revealed a significant shift of *Kmt2a* isoforms in the *Kmt2a^+/LSL^* samples, from the full length canonical *Kmt2a-201* isoform to the truncated *Kmt2a-203* isoform, that does not contain the enzymatic SET domain (Figure 5G-H). More notably, upon the LSL cassette removal in the Nes-Cre^+/-^*Kmt2a^+/LSL^* cells, these effects are reversed, confirming functionality of the cassette. Taken together, our findings support the notion that a heterozygous loss of *Kmt2a* leads to aberrant mRNA expression in mNPCs. These changes at the molecular level are then reversed *in-utero* in the Nes-Cre^+/-^*Kmt2a^+/LSL^* model.

### H3K4me1 signal is rescued but does not drive gene expression changes in Nes-Cre^+/-^*Kmt2a^+/LSL^* mNPCs

To look further into the effects of heterozygous *Kmt2a* loss at the chromatin level, we performed a quantification of the H3K4me1 mark using CUT&RUN in mNPCs (n=3 *Kmt2a^+/+^*, n=4 *Kmt2a^+/LSL^* and Nes-Cre^+/-^*Kmt2a^+/LSL^*). When comparing *Kmt2a^+/+^* and *Kmt2a^+/LSL^* samples, the majority of differential peaks (70.4%, 1380/1962 peaks) show a gain of H3K4me1 in *Kmt2a^+/LSL^* samples (Figure 4A, Supplemental Table 3). While mapping enhancer regions to genes is not trivial, when we linked the disrupted peaks to the nearest gene by distance, we found no correlation between the gene expression changes and the H3K4me1 epigenetic mark (Figure 4B-C, See Methods). Investigating the increased H3K4me1 signal further, we compared the signal over mouse NPC enhancer and neural enhancer regions from EnhancerAtlas [32]. This revealed an increased signal at the neuronal enhancers but not at the NPC enhancers for *Kmt2a^+/LSL^* samples (Figure 4D). Again, the increased signal does not correlate with the nearest gene’s expression changes (Supplemental Figure 6A). To assess the rescue of this epigenetic mark in Nes-Cre^+/-^*Kmt2a^+/LSL^* samples, we observe a reversal of 98.0% of peaks (Supplemental Figure 6B) and when visualizing the overall H3K4me1 signal at the differential peaks on a heatmap, we observe an increased signal in *Kmt2a^+/LSL^* samples compared to *Kmt2a^+/+^* wild-types, which is then reduced in Nes-Cre^+/-^*Kmt2a^+/LSL^* samples (Figure 4E). This effect can be more specifically visualized on tracks showing gained H3K4me1 signal at neuronal enhancers for genes *Acsl3* and *Bahcc1* for *Kmt2a^+/LSL^* samples (Figure 4F). When applying a more relaxed cut-off, the 4963 peaks that fell into the top decile of p-values when overlapping the results from *Kmt2a^+/LSL^* vs *Kmt2a^+/+^* samples and the *Kmt2a^+/LSL^* and Nes-Cre^+/-^*Kmt2a^+/LSL^* samples, map to 3646 individual genes. When performing an over-representation analysis (ORA) of these genes, we observe pathways involved in CNS differentiation, development, and neurogenesis (Figure 4G), highlighting that the disruption is occurring at functionally relevant sites. When examining the genes with restored H3K4me1 signal and gene expression, we observe a non-significant overlap of genes with a gained H3K4me1 peak and increased expression in *Kmt2a^+/LSL^* samples (Figure 4H), as well as a non-significant overlap of genes with a lost H3K4me1 peak and decreased expression in *Kmt2a^+/LSL^* samples (Figure 4I). Collectively, we observe an increased H3K4me1 signal in *Kmt2a^+/LSL^* samples, albeit with no global effects on gene expression.

**Figure 4:**
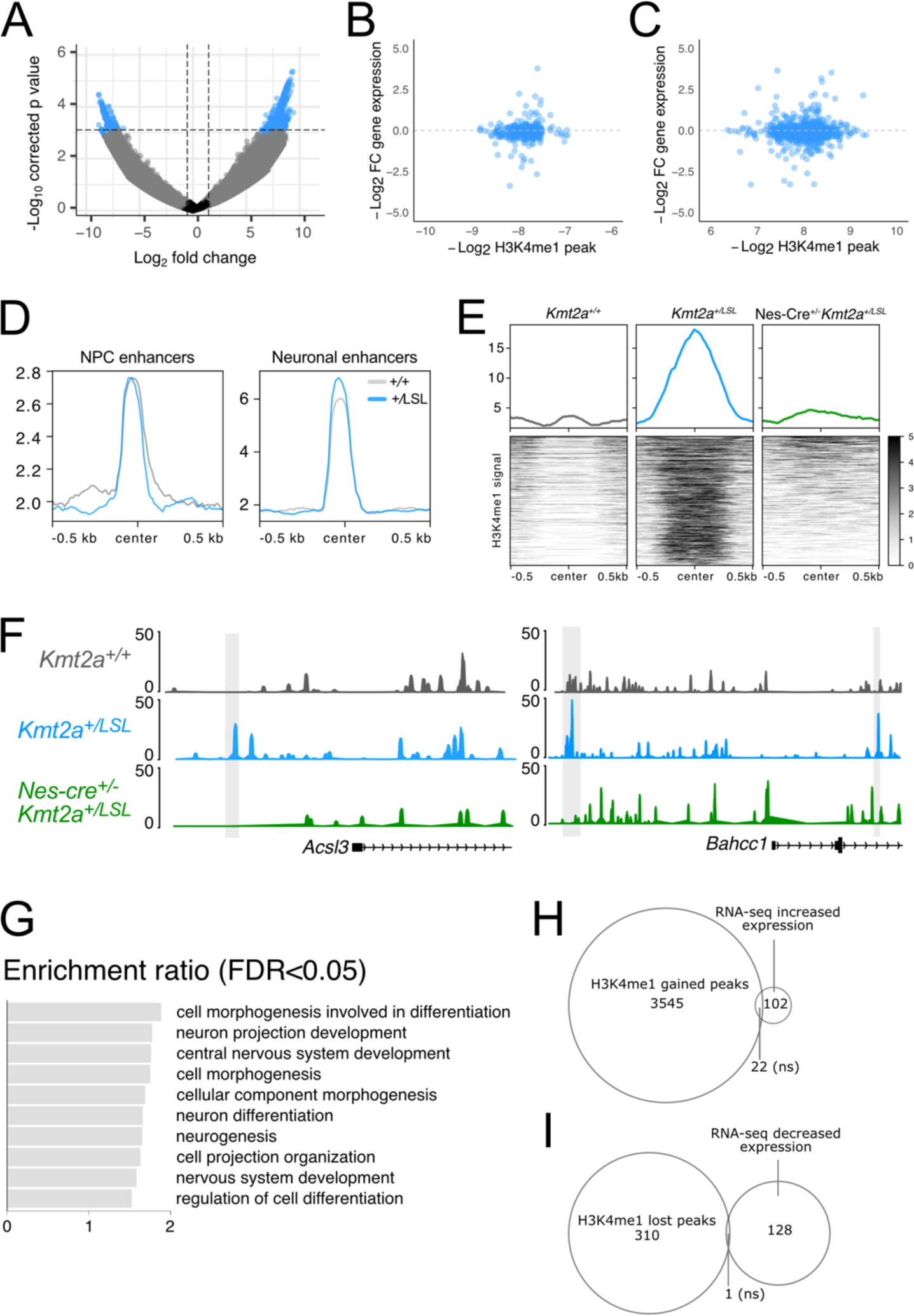
Epigenetic imbalance caused by heterozygous *Kmt2a* loss. (A) Volcano plot showing differential H3K4me1 peaks comparing *Kmt2a^+/+^* and *Kmt2a^+/LSL^* samples. Expression changes for genes with (B) lost H3K4me1 peaks and (C) gained H3K4me1 peaks in *Kmt2a^+/LSL^* samples. (D) Signal plot of H3K4me1 signal for *Kmt2a^+/+^* and *Kmt2a^+/LSL^* samples over NPC and neuronal enhancers. (E) Heatmap of H3K4me1 signal at differential sites when comparing *Kmt2a^+/+^*, *Kmt2a^+/LSL^* and Nes-Cre^+/-^*Kmt2a^+/LSL^* samples. (F) Representative H3K4me1 signal figures, showing gained signal in *Kmt2a^+/LSL^* samples at neuronal enhancers (gray) at *Acsl3* and *Bahcc1* genes compared to wild-type and rescued samples. (G) Over-representation analysis of genes with differential H3K4me1 peak in *Kmt2a^+/LSL^*samples. Image shows enriched pathways (with FDR<0.05). (H) Overlap of genes with a gained H3K4me1 peak and genes that have an increased expression in *Kmt2a^+/LSL^* samples. (I) Overlap of genes with a lost H3K4me1 peak and genes that have decreased expression in *Kmt2a^+/LSL^* samples. ns: not significant, *p< 0.05, **p < 0.01, ***p < 0.001, ****p < 0.0001.

**Figure 5:**
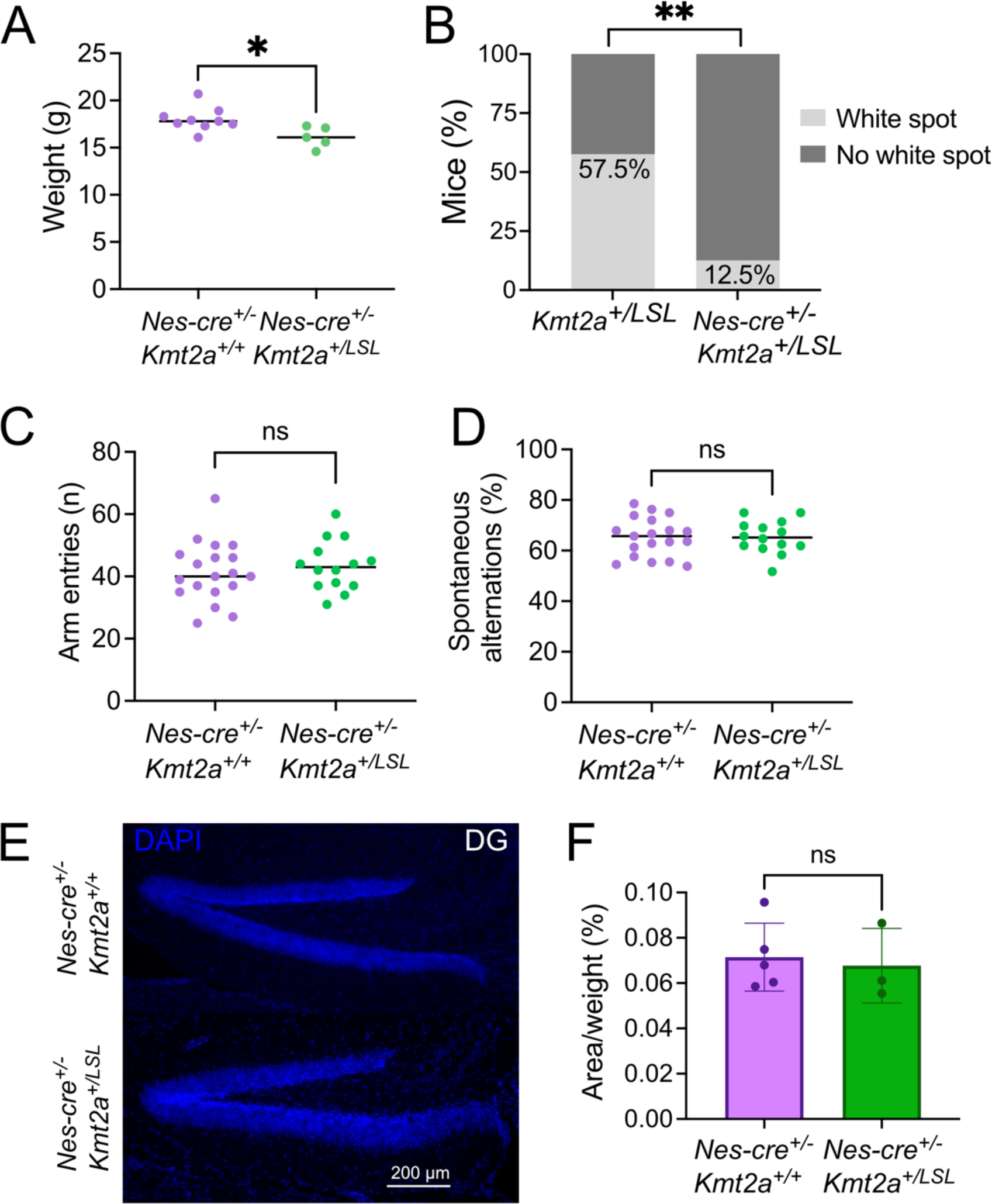
*In-utero* rescue of the neurological phenotypes in WDSTS mice. (A) The Nes-Cre^+/-^*Kmt2a^+/LSL^*mice demonstrate growth deficiency compared to their Nes-Cre^+/-^ *Kmt2a^+/+^*littermates (n=5-9). (B) The presence of the white belly spot is significantly reduced in Nes-Cre^+/-^*Kmt2a^+/LSL^* mice (12.5%) compared to *Kmt2a^+/LSL^* mice (57.5%) (Fisher’s exact test). (C) Nes-Cre^+/-^ *Kmt2a^+/LSL^* mice do not show a significant decrease of total arm entries in a Y maze (n=14-19 per genotype) (D) Nes-Cre^+/-^*Kmt2a^+/LSL^* mice do not exhibit differences in spontaneous alternations in the Y-maze, compared to their Nes-Cre^+/-^*Kmt2a^+/+^* littermates. (E) Representative images of DG of the hippocampus area in Nes-Cre^+/-^*Kmt2a^+/+^* and Nes-Cre^+/-^*Kmt2a ^+/LSL^* mice. (F) Quantification of granule cell layer area of the DG of the hippocampus demonstrating no difference in size when corrected for body weight (n=3-5 per genotype). ns: not significant, *p< 0.05, **p < 0.01, ***p < 0.001, ****p < 0.0001.

### WDSTS phenotypes are rescued *in-utero*

After demonstrating that nestin-Cre leads to sufficient recombination of the LSL cassette and rescued *Kmt2a* expression (see Figure 3, Supplemental Figure 4) we characterized the Nes-Cre^+/-^*Kmt2a^+/LSL^* mice. The Nes-Cre^+/-^*Kmt2a^+/LSL^* mice demonstrate growth deficiency compared to their Nes-Cre^+/-^*Kmt2a^+/+^* littermates (n=5-9, Figure 5A, Supplemental figure 7A-B). Furthermore, we found similar craniofacial defects of the Nes-Cre^+/-^*Kmt2a^+/LSL^* as previously described in *Kmt2a^+/LSL^* mice (Figure 1E-H) indicating that there is no rescue of this phenotype (Supplemental figure 7C-D). This would suggest that either the growth defects of the *Kmt2a^+/LSL^* mice are not solely driven by nestin-expressing cell types, or that the cause of the growth phenotypes occurs prior to when the rescue of *Kmt2a* takes place *in-utero*. In contrast, the prevalence of the white abdominal spot of the Nes-Cre^+/-^*Kmt2a^+/LSL^* mice drastically decreased from 57.5% to 12.5% (Figure 5B) compared to the 57.5% of the *Kmt2a^+/LSL^* mice, indicating malleability of this phenotype. Behavioral testing of adult mice demonstrates no difference of arm entries between Nes-Cre^+/-^*Kmt2a^+/+^* and Nes-Cre^+/-^ *Kmt2a^+/LSL^* (n=14-19, Figure 5C), nor of the percentage of spontaneous alterations in the Y-maze (Figure 5D), indicative of non-impaired working memory of Nes-Cre^+/-^*Kmt2a^+/LSL^* mice. Similarly, the brain histology of the Nes-Cre^+/-^*Kmt2a^+/LSL^* mice demonstrate normal hippocampal structure, with a non-significant difference in the size of the area of the granule layer of the DG, when compared to Nes-Cre^+/-^*Kmt2a^+/+^* littermates and corrected for body weight (Figure 5E-F). Taken together, when bred to a nestin-Cre mouse model, we observe histological and behavioral rescue of the neurological defects of the Nes-Cre^+/-^*Kmt2a^+/LSL^* mice.

## Discussion

In this paper we have created and characterized a novel WDSTS mouse model (*Kmt2a^+/LSL^).* This model depicts many of the human WDSTS phenotypes, validating it as a disease model for WDSTS. We observed rescue of the neurological phenotype of the mouse model after crossing the mice to a nestin-Cre model, demonstrating *in-utero* malleability of WDSTS. Genetic disorders are being diagnosed earlier than before with the onset of whole genome sequencing and exome sequencing [33] and there is an explosion of strategies that have successfully made it to the clinic including small molecules, ASO-based strategies and CRISPR-Cas9 based strategies [34], [35]. Using a genetically based strategy to examine the likelihood, the window of opportunity, and which aspects of the phenotype are malleable could be of great value for prioritizing any Mendelian disorder for therapeutic development as well as offering an excellent genetic control for such studies. Our work uses one such strategy which could be used for other congenital disorders to test the timing and effectiveness of early manipulation. We demonstrate rescue of the altered *Kmt2a* allele and its consequences on multiple levels: genetic, molecular, and phenotypic. Our work also provides temporal insights into the timing of disease onset in the CNS and neural crest. There is some debate in the literature surrounding the exact timing of action in the nestin-Cre mouse model, with the original paper suggesting recombination occurring as early as E11 in the nervous system [36], [37]. However, more recent studies have shown that Cre expression only reaches sufficiently high levels in NSCs and NPCs to cause recombination during late embryonic (E15.5-17.5) and early postnatal periods [36].

*In vitro*, we find robust evidence that the transcriptional changes caused by the loss of *Kmt2a* are reversed, with the majority of genes downregulated upon its loss and then restored. Surprisingly, the directionality of the chromatin-rescue occurs in the opposite direction from what would be expected for an H3K4 methyltransferase, i.e. the sites that show the biggest effect have increased levels of the H3K4me1 mark for *Kmt2a^+/LSL^* samples, that then normalizes upon genetic rescue. A similar chromatin effect, with a trend towards increased chromatin accessibility, has previously been observed in studies of patients and animal models of Kabuki syndromes [38], [39] and in cells from animal models of Rubinstein-Taybi and Kleefstra syndromes [7], [40]. Moreover, the increased H3K4me1 signal at neuronal enhancers could suggest the cells are priming these enhancers towards a premature neuronal phenotype, similar to what has been described in Kabuki syndrome [41], [42]. Another potential reason for this observation could be that the increased H3K4 methylation involves a compensatory response. For instance, KMT2A might act as a direct or indirect negative regulator of another epigenetic regulators that promote open chromatin. We also observe signs of increased transcriptional initiation of *Kmt2a* and isoform switching in *Kmt2a^+/LSL^* samples, showing attempted compensation is taking place at least at the locus itself. Alternatively, we hypothesize KMT2A might be occupying binding sites, effectively blocking other epigenetic-or transcriptional factors from binding at those loci. We also find no correlation of gene expression linked to the changes in H3K4me1 signal, suggesting that the lack of KMT2A’s enzymatic function is not the sole driver of WDSTS pathogenesis. This aligns with our prior finding that WDSTS missense variants do not show enrichment within the enzymatic SET domain of KMT2A in contrast to e.g. Kabuki syndrome causing missense variants that cluster in the KMT2D-SET domain [16][43]. This is emphasized even further with the fact that enzyme-specific homozygous KO of *Kmt2a* yields viable and fertile mice while such KOs lead to lethality in other *Kmt2* mouse models e.g. *Kmt2d* [11][44]. Further studies will be needed to understand the underlying mechanism of this effect in more detail. Our findings suggest that disruption of gene expression and a global imbalance in the epigenetic machinery are not solely caused by the loss of the enzymatic activity of KMT2A and may result in cellular compensation, leading to downstream secondary effects, which could play a major role in MDEM disease mechanisms.

In addition to previously described features, the *Kmt2a^+/LSL^* model demonstrates multiple additional common WDSTS-like phenotypes such as craniofacial defects, hypotonia and hypertrichosis, one of the defining features of WDSTS (Supplemental Table 4). Interestingly, our mice appear to show increased hair follicle density in some regions, without an obvious rate of regrowth of the hair, and could therefore open a novel avenue of research into the mechanistic basis of hypertrichosis in humans. We observe no rescue of the growth retardation of Nes-Cre^+/-^*Kmt2a^+/LSL^* mice. However, it is important to note that the nestin-Cre mice have previously been shown to demonstrate mild hypopituitarism, accompanied by decreased body weight, and to ensure that these results could not be due to the nestin-Cre genotype, we compared their weight to the Nes-Cre^+/-^*Kmt2a^+/+^* littermates [45]. Similarly, we observed no rescue of the craniofacial structure abnormalities in the Nes-Cre^+/-^*Kmt2a^+/LSL^* mice. This could be explained in two ways. Either the growth retardation might be related to dysfunction in cell populations that do not express nestin, or secondly, the growth retardation might occur early enough in development to not be rescued by the nestin-Cre expression. In both scenarios we would not expect a rescue of these phenotypes.

Previous studies have linked abnormal craniofacial features to neural crest dysfunction in KS in mice [19], [20], [21], a prominent phenotype shared between KS and WDSTS. In concordance, we observe a prominent white abdominal spot in the majority of *Kmt2a^+/LSL^* mice, with a striking decrease in the Nes-Cre^+/-^*Kmt2a^+/LSL^* mice, without fully eliminating it. This could suggest the timing of rescue of the neural crest defects is around the same time as nestin expression starts, showing partial rescue in our model. Ultimately, our findings suggest a role for *Kmt2a* in neural crest development with more research needed to fully elucidate its role. Lastly, we observe rescue of the neurological dysfunction in the Nes-Cre^+/-^*Kmt2a^+/LSL^* mice both histologically and in behavior. We conclude the neurological phenotype of WDSTS is malleable *in-utero* and that WDSTS is yet another treatable cause of ID from the group of Mendelian disorders of the epigenetic machinery.

## Acknowledgements

We thank the Wiedemann-Steiner foundation for supporting this work. We thank Leandros Boukas, Katrín Möller and Hilmar Örn Gunnlaugsson for valuable input. Some figures were created using BioRender.com. Data utilized in this publication was provided by RARE-X, a research program of Global Genes. RARE-X provides a collaborative platform for global data sharing and analysis to accelerate research for rare diseases. Global Genes is a 501(c)(3) non-profit organization dedicated to eliminating the burdens and challenges of rare diseases for patients and families globally.

## Funding

This work was specifically funded by a grant to HTB from the Wiedemann-Steiner Foundation. HTB and JO are also funded by a grant from the Icelandic research fund (217988-051). TR and JO are funded by individual grants from the University of Iceland doctoral fund grants. Mouse model strain development, strain curation and cryopreservation of the mouse strains was supported by the Jackson Laboratory Precision Genetics Grant from the National Institutes of Health, U540D030157 to C.M.L.

## Author contributions

Conceptualization: HTB; Methodology: HTB, TR, KJA, JO, AOS, VBD, ARZ; Validation: TR, KJA, ARZ; Formal analysis: TR, KJA, VBD, AB, KPF; Investigation: TR, KJA, AOS, JO, ARZ; Resources: HTB; Data curation: TR, KJA, LB, VBD, AB, KPF; Writing – original draft: TR; Writing – review & editing: HTB, TR, KJA; Visualization: TR, KJA, VBD; Supervision: HTB; Project administration: TR, HTB, KJA; Funding acquisition: HTB, CML.

## Competing interests

HTB is a consultant for Mahzi Therapeutics and founder of Kaldur Therapeutics. AOS is an employee of Arctic Therapeutics. Other authors declare that they have no competing interests.

## Methods

### Animals

The mouse model was created by the Jackson Laboratory, by microinjection of C57BL/6J single cell zygotes with CRISPR/Cas9 using guides 5’-CTGCAGCGAGAGACTGTATG-3’ and 5’GAGACTGTATGAGGTATCAG-3’ with a donor plasmid containing a lox-STOP-lox element flanked by *Kmt2a* intron 1 genomic DNA sequences of 1.4-kb and 1.8-kb, respectively. Of 38 mice recovered, six founders were identified with the desired genome edited allele at the Kmt2a locus, as determined by genomic long-range PCR. Two lines were established after mating of the founders to C57BL/6J mice to confirm germ line transmission of the edited lox-STOP-lox knock-in (KI) allele in *Kmt2a* and subsequently expanded by an additional backcross to C57BL/6J mice (JAX Stock #664) prior to intercrossing and strain validation [46]. These two conditional KI (cKI) lines, designated as stock numbers #35724 C57BL/6J-*Kmt2a^em7Lutzy^*/J and #37525 C57BL/6J-*Kmt2a^em8Lutzy^*/J each contain the same allele but were derived from different founders. The functionality of the loxP elements within each KI allele was confirmed by test mating of heterozygous #35724 and #35725 male mice with hemizygous B6.Cg-*Edil3^Tg(Sox2-cre)1Amc^*/J female Cre-expressing mice (JAX Stock #8454) [47]. Whereas both strains were mated with Sox2-Cre to confirm loxP functions, postCre Het x Het matings were only performed using the em7.1 allele (JAX Stock #35727, now extinct). Sox2-Cre Tg did not segregate in these mice (Supplemental Figure 1G). This validation revealed both stocks #35724 and #37525 behaved the same way in subsequent matings and therefore only Stock #35724, named *Kmt2a^+/LSL^*, was characterized further with respect to expression profiling in the pre-and post Cre mice. Jackson laboratory Stock #35724 can be purchased from the Jackson Laboratory mouse mutant repository (https://mice.jax.org). Genotyping was performed from DNA isolated from ear clips using primers Kmt2a-genoF1-(5’-ACA CAT GGC TTC CTG GAG G-3’) and Kmt2a-genoR1-(5’-CTG TTT GCA GTC GGA AAG CC-3’). Nestin-Cre^+/+^ mice (JAX stock #3771) were acquired from the Jackson Laboratory [48], [49]. Nestin-Cre model was genotyped using primers NesCre-WT-F-(5’-TTG CTA AAG CGC TAC ATA GGA -3’), NesCre-Tg-F-(5’-CCT TCC TGA AGC AGT AGA GCA-3’), NesCre-R-(5’-GCC TTA TTG TGG AAG GAC TG -3’). All mouse strains were maintained on pure C57BL/6J background and provided food (Altromin NIH#31 M (breeding) or Altromin 1324 (maintenance), Brogaarden) and filtered water *ad libitum*. The animals were maintained under a 12-hour light/dark cycle at room temperature (21-23°C) with relative humidity around 40%. Regular FELASA health monitoring was conducted with samples taken from sentinel and resident animals and analyzed by IDEXX Germany, yielding negative results during experimental period. Import of mice and experiment protocols were in accordance with Icelandic Food and Veterinary Authority (license no. 2208602) and accepted by National Expert Advisory Board on Animal Welfare.

### Behavioral testing

The hypotonia tests (surface righting and hindlimb suspension tests) were performed on P6 pups as described by Feather-Schussler et al [50]. For the surface righting test, the pups were placed on their backs and held in place for 5 seconds, recording the time for the pups to return to a prone position. The test was conducted in 3 trials, giving the pup a 30 second rest in between each trial and a time-out at 60 seconds, with mean scores over all trials plotted. For the hindlimb suspension test, the pups were placed on their hindlimbs on the rim of a 50 ml conical tube, with the latency to fall recorded. The test was conducted in 3 trials, giving the pup a 60 second rest in between each trial and a time-out at 5 minutes. All trials were plotted, with each trial tiring the mice more. The spontaneous alternation Y-maze was performed on 8–10-week-old mice according to protocol described by Kraeuter et al [51]. Spontaneous alternation percentage is calculated with the following formula:

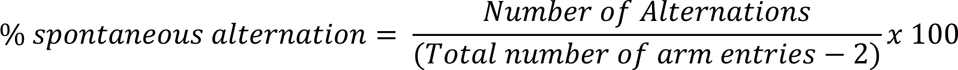

### CT scan

CT scans were performed on 2-3.5-month-old mice (3-6 mice per genotype) using 50 kV tube voltage, 0.21 mA tube current and 1712 mGy dose estimate using the pre-clinical optical imaging and CT system from MILabs (Houten, Netherlands). Virtual reconstructions of the skull were produced for each specimen by importing the CT data to 3dSlicer at half resolution (0.030 x 0.030 x 0.030 mm^3^ cubic voxels), and generating a model of the skull surface based on thresholding. A geometric morphometric approach allowed us to estimate cranial *shape* based on 3D coordinate data for a set of 31 biologically homologous landmarks (Supplemental Table 5) evenly distributed across the cranium. Landmark coordinate data were exported from 3dSlicer [52] and we used the R *geomorph* package (v.4.0.7) for all morphometric analyses [53]. We assumed object symmetry for each specimen using the *bilat.symmetry*() function, which removes the effect of asymmetric variation [54]. For a subset of our data, rostral and caudal ends of the crania were inadvertently truncated in the original CT scanning episode. This resulted in missing data for nasale (nal) in six specimens, and for opisthion (opi), basion (bas), and occipital condyles (rfmc and lfmc) in 11 specimens. Missing data were estimated using the *estimate.missing()* function, which predicts the location of those landmarks based on the average of other individuals in the sample [55]. Both *Kmt2a^+/LSL^* and wild-type genotypes were equally represented in this average sample. For each analysis, we conducted a new Procrustes superimposition and tested the effect of genotype using a Procrustes ANOVA with the *procD.lm()* function. We visualized the results using Principal Components Analysis (PCA).

### Histology

Samples were taken from the belly and backs of newborn pups, fixed in 4% paraformaldehyde, and then processed at the Department of Pathology, Landspitali National University Hospital, Reykjavik, Iceland. The tissue samples were paraffin-embedded and cut into 1.5µm sections for hematoxylin and eosin (H&E) staining and 3µm sections for immunohistochemistry (IHC). H&E staining was done using standard methods. Sections for immunohistochemistry were de-paraffinized and rehydrated in xylene and ethanol. Antigen retrieval was done with Envision Flex Target Retrieval Solution, high 9 (Agilent, K800421-2). Sections were immunostained in AutostainerLink 48 from Agilent with EnVision Detection System Peroxidase/DAB, Rabbit/Mouse kit (Agilent, K4065). Incubation with primary antibody Calretinin (Agilent, M7245) was performed at room temperature for 30 min. After incubation with a primary antibody, sections were incubated with EnVision FLEX/HRP. Sections were washed between steps with Tris-buffered NaCl solution with Polysorbate 20 pH 7.6 (Agilent, S3306). All sections were incubated with 3,3ʹ-diaminobenzidine solution (Agilent, K4065) for 10 min. Sections were counterstained with hematoxylin for 30 seconds. Finally, sections were dehydrated with 100% ethanol and xylene followed by cover slipping with mounting medium (Pertex, Histolab). Images were acquired with NanoZoomer XR (HAMAMATSU) scanner. Sections were analyzed using NDP view (version 2.7.25). Hair follicles were counted on 8 mm of skin, each section was counted 4 times, and the average number was plotted per mm of skin.

### CBC measurement and hair removal

Blood samples for Complete Blood Count (CBC) were collected in EDTA blood tubes from 3-5 m/o mice (n=8 per genotype) and measured on VetScan HM5 v2.1 (Abaxis Inc, USA). Hair was removed from the backs of adult mice using Veet Hair Removal Crème (Reckitt Benckiser).

### Perfusion, cryosectioning, IF staining

Mice were euthanized 9 weeks old with Xylasine (4 mg/ml final concentration) Ketamine (16 mg/ml concentration) combination in 0.9% NaCl and perfused intravascularly with 4% PFA after first flushing blood system with 1x PBS. Brains were dissected out and cryopreserved in a two-step gradient of 30% and 15% sucrose solution (sucrose in 0.1M phosphate buffer pH=7.2). Brains were cut sagittally and frozen in Tissue-Tek OCT compound. Brains were sectioned in 20µm serial sections on Leica CM1850 cryostat (Leica Biosystems). Sections were stored at -80°C until stained. Every 2^nd^, 6^th^, 10^th^, and 15^th^ brain section were used for DAPI staining. Samples were washed 2×10 minutes with TBS-T (1x TBS (Santa Cruz, sc-24951) with 0.05% Triton X-100 (Sigma-Aldrich, T8787-100ML)) and then treated in DAKO (sodium citrate buffer pH=6) at 95°C for 20 minutes for antigen retrieval, washed 3×10 minutes in TBS-T and then blocked for an hour in TBS-T+ (TBS-T with 3% normal goat serum (Abcam, ab7481)) at room temperature. Samples were incubated in primary antibody solution (antibody in TBS-T+) overnight at 4°C. Slides were washed 3x 10 minutes in TBST-T and mounted with Fluoromount-G with DAPI (Thermo Fisher Scientific, 00-4959-52). Primary antibodies used Alexa Fluor® 647 Anti-NeuN antibody (Abcam, ab190565 1:500 dilution). Brain samples were imaged using CrestOptics CICERO spinning disk with a 10x and 40x objective. Images were processed using Fiji [56]. DAPI staining were used for area quantification, representative NeuN stainings in Supplemental Figure 2F.

RNA isolation from brain and Real Time Quantitative Polymerase Chain Reaction (RT-qPCR) Total RNA was isolated from whole brain tissue of P1 pups using TRIzol™ Reagent (Thermo Fisher Scientific, #15596026) according to the manufacturer’s instructions. RNA concentration was measured on NanoDrop (Thermo Fisher Scientific), and 0.5 ng of RNA was used for input for cDNA synthesis using High-Capacity cDNA Reverse Transcription Kit (Applied Biosystems, #4368814), run on the MiniAmp Thermal Cycler (Applied Biosystems). Luna Universal qPCR Master Mix (NEB, M3003) was used for RT-qPCR reactions and run on CFX384 Real-Time PCR Detection System (Bio-Rad). Each biological replicate of the RT-qPCR assay in this study was carried out in technical triplicates.

### Mouse NPC extraction and culture

Mouse NPCs were dissected from hippocampus of P0 pups and cultured in an adapted protocol from Bernas et al [57]. Dissected hippocampi were dissociated in 1X TrypLE™ (Thermo Fisher Scientific, A1217701), washed in Neurobasal growth medium: Neurobasal medium (Thermo Fisher Scientific, 21103049) with 1X B27 supplement (Thermo Fisher Scientific, 17504044) and filtered through a 70 µm cell strainer (Miltenyi Biotech, 130-110-916). The cell suspension was centrifuged at 350g for 5 minutes, resuspended in 500µl Neurobasal growth medium containing 1X Penicillin/Streptomycin (Sigma, P4333-100ML), 1X Glutamax (Thermo Fisher Scientific, 35050038), 20ng/mL FGF-2 (Peprotech, 100-18B), 20 ng/mL EFG (Peprotech, AF-100-15), and 2 µg/mL heparin (MP Biomedicals, 210193125) and seeded onto Matrigel (Corning, 354234) coated plates that were previously incubated at 37°C for 2 hours (Matrigel diluted 1:100 in DMEM/F-12 with GlutaMAX™ Supplement (Thermo Fisher Scientific, 31331028)). After first splitting of cells, mNPCs were cultured in media conditions listed above on plates coated firstly with poly-D-Lysine (PDL) hydrobromide (Sigma, P7280) overnight followed by laminin (Roche, 11243217001) coating for 2 hours at 37°C.

### RNA-seq

mNPCs were harvested in TRIzol™ Reagent (Thermo Fisher Scientific, #15596026) and RNA isolated with Direct-zol™ RNA Microprep kit (Zymo, #R2060). RNA concentration and purity was measured on a Nanodrop and with Agilent RNA 6000 Nano Kit (Agilent Technologies, 5067-1511) on Agilent 2100 Bioanalyzer. All samples had RIN>9. Samples were sequenced by sequencing company Novogene (Cambridge, UK). Raw RNA-seq data (fastq files) were downloaded from Novogene and raw reads were pseudoaligned to the mouse mm10 reference transcriptome (GRCm38) using Kallisto (version 0.48.0) [58], running the kallisto quant command with-b 100 for 100 bootstraps. For viewing sequencing coverage, fastq files were aligned to mm10 reference transcriptome using kallisto quant using-genomebam, creating bam files that were converted to bigWig files using bamCoverage with default settings from deepTools2 (version 3.5.1) [59]. Genome coverage bigwig files were viewed in the UCSC browser [60]. Aligned data was imported to R [61] using tximport (version 1.30.0) [62] and mapped to mm10 genes with the BiomaRt package [63]. Differential expression analysis was performed on samples containing >10 counts and genes with only one sample containing counts were excluded. We used DESeq2 (version 1.42.0) using default settings [64]. To correct for false discovery rates, the z-scores returned from DESeq2 were used as an input for fdrtool with statistic=”normal” and statistical cut of was set at corrected q-value < 0.05 [65]. For further analysis we used three genotypes: *Kmt2a^+/+^* for wildtype samples, *Kmt2a^+/LSL^* for WDSTS samples and Nes-Cre^+/-^*Kmt2a^+/LSL^* for assessing the *in-utero* rescue, excluding the Nes-Cre^+/-^*Kmt2a^+/+^* cells as they served only as another wildtype control. To assess the quality of the differential analysis we performed three quality control checks. First, we inspected the histogram of corrected p-values (Supplemental Figure 5D), revealing an even distribution of values between 0 and 1, with a clustering of values close to 0. This suggests a calibrated differential analysis. Secondly, we observed the genotype differences on a principal component analysis (PCA) plot (Supplemental Figure 5E), after performing regularized log transformation with rlog() function on expression matrix. PCA plot reveals slight variance between biological replicates within genotypes. Lastly, an MA plot demonstrates no obvious systematic issues of the data (Supplementary Figure 5F). WebGestalt was used for over-representation analysis (ORA) [66]. Autism-spectrum-disease (ASD) risk genes were acquired from SFARI Gene database (https://gene.sfari.org/, version Q1 2024) [31]. The Complex Heatmaps package was used for generating heatmaps for gene expression visualization (version 2.18.0) [67]. Expression values were normalized to z-scores and outliers with z score>2 were filtered out, with the rest plotted. IsoformSwitchAnalyzeR was used for identification of isoform switches (version 2.2.0) [68]. Venn diagrams were created with DeepVenn [69] and Fisher’s test was applied for statistical analysis using GeneOverlap [70].

### CUT&RUN

CUT&RUN was performed according to CUTANA™ CUT&RUN Protocol (v.1.7 EpiCypher) from 160.000 cells per sample, with n=3 *Kmt2a^+/+^*, n=4 *Kmt2a^+/LSL^* and n=4 Nes-Cre^+/-^*Kmt2a^+/LSL^*. Cells were permeabilized with 0.01% digitonin buffer (Sigma, D141) and E. Coli DNA (EpiCypher, 23618-1401) was used as spike-in for normalizaÄon (0.2 ng final concentraÄon per sample). The anÄbodies used were Rabbit IgG AnÄbody (EpiCypher, 23613-0042, 0.5µl per sample) and H3K4me1 (Abcam, ab8895, 0.25µl per sample). DNA was isolated using Genomic DNA Clean & Concentrator (Zymo, D4065). Library preparaÄon was done using TruSeq RNA CD Index plate (Illumina, 20019792). Quality of samples were assessed using Agilent Bioanalyzer DNA high sensiÄvity kit (Agilent Technologies, 5067-4626). DNA samples were pooled to a 1.8pM concentration and sequenced on a NextSeq 550Dx instrument (Illumina) using NextSeq 500/550 High output kit v2.5 (Illumina, 20024907). Demultiplexing was performed using Illumina bcl2fastq command with the options--ignore-missing-control--ignore-missing-positions--ignore-missing-filter. Fastq sample files were aligned to the mouse mm10 genome using Bowtie2 (version 2.4.4) [71] for paired end reading. Aligned sam files were sorted, indexed and converted to bam files using SAMtools (version 1.17) [72]. For visualizing H3K4me1 peaks in UCSC, bam files for all samples for each genotype were merged with SAMtools. They were then converted to bigwig files with deepTools (version 3.5.1) and normalized to 1x mouse genome with--normalizeUsing RPGC [59]. Peaks were called for merged genotypes against the merged IgG samples for each genotype using MACS3 (version 3.0.0) with settings-q 0.1,-maxgap-300,-keep-dup all [73]. First, the raw reads for each sample were counted using dba.count() function from Diffbind (version 3.12.0) with „score= DBA_SCORE_READS“, followed up with retrieving the matrix with dba.peakset() function, with settings „bRetrieve=TRUE“ [74]. Raw count matrix was filtered to only include peaks where the was greater than 10 for at least one genotype. Count matrix was used as input for DESeq2 where differential analysis was performed. FDRtool with statistic=”normal” was applied to correct for false discovery rates, with a cut off at corrected p-value <0.001. H3K4me1 peaks were annotated to nearest gene using annotatePeak() function from ChIPseeker package (version 1.5.1) [75]. Mouse NPC and Neuronal enhancer sites were retrieved from Enhancer Atlas (version 2.0) [32]. Heatmaps and plotprofiles were plotted using deepTools2 (version 3.5.1) [59].

### GWS

DNA from mNPCs in culture were harvested from *Kmt2a^+/LSL^* cell line cell line (1 million cells) using Monarch® HMW DNA Extraction Kit (NEB, T3050). We sheared the high molecular weight (HMW) DNA to achieve consistent and narrow size ranges to 20kb using the Megaruptor 3 Shearing kit (E07010003, Diagenode). Megaruptor 3 was performed using customized settings, including a concentration of 50ng/µL, a volume of 70µL, and a speed of 30. Subsequently, we measured the sheared DNA size using the HS Large Fragment 50kb kit (DNF-464-0500, Agilent) on a Fragment Analyzer and analyzed by the High Sensitivity Large Fragment 50 kb Analysis software. We performed library preparation with a Ligation Sequencing kit V14 (SQK-LSK114, Oxford Nanopore Technologies, ONT), and sequenced by loading the sample into an R10.4.1 flow cell (FLO-PRO114M, ONT) loaded on a PromethION 24 sequencer based on the manufacture’s guidelines (ONT). Nanopore sequencing was conducted using customized run settings, which included a Run Limit of 96 hours, a Minimum Read Length of 200 bp, and Super-accurate Base calling. To optimize output, we washed the flow cell once after 22 hours of sequencing using the Flow Cell Wash kit XL (EXP-WAS004-XL, ONT), followed by a 1.5-hour wait period and flushing of the flow cell with Flow Cell Priming Mix twice by using Sequencing Auxiliary Vials V14 (EXP-AUX003, ONT), We utilized the remaining DNA library from the previous preparation for loading. Data acquisition and base calling in real-time using PromethION device and MinKNOW software developed by ONT. Nanopore sequencing yielded approximately 86.43 Gb of estimated bases with an average read length of 23.5 kb. All fastq files were merged into one fastq file using cat command in Linux. A sam file was created by aligning the fastq file to the mouse genome using minimap2 with –map-ont function for Oxford Nanopore Technologies data [76] and reference genome mm10. Sam file was converted to a bam file, sorted and indexed using samtools [72]. Sequencing results were viewed in IGV [77].

### Statistical testing

Data was analyzed using Prism 9 (v 9.4.1, GraphPad Software, Boston Massachusetts USA) and R [61]. Unless otherwise stated, significance between two groups was calculated with an unpaired t-test using a P-value nomenclature; *p< 0.05, **p < 0.01, ***p < 0.001, ****p < 0.0001.

## References

[1] W. D. Jones et al., “De novo mutations in MLL cause Wiedemann-Steiner syndrome,” Am. J. Hum. Genet., vol. 91, no. 2, pp. 358–364, Aug. 2012, doi: 10.1016/j.ajhg.2012.06.008.

[2] T. A. Milne et al., “MLL targets SET domain methyltransferase activity to Hox gene promoters,” Mol. Cell, vol. 10, no. 5, pp. 1107–1117, Nov. 2002, doi: 10.1016/s1097-2765(02)00741-4.

[3] T. Kusch, “Histone H3 lysine 4 methylation revisited,” Transcription, vol. 3, no. 6, pp. 310–314, 2012, doi: 10.4161/trns.21911.

[4] H. Santos-Rosa et al., “Active genes are tri-methylated at K4 of histone H3,” Nature, vol. 419, no. 6905, pp. 407–411, Sep. 2002, doi: 10.1038/nature01080.

[5] Y.-C. Huang et al., “The epigenetic factor Kmt2a/Mll1 regulates neural progenitor proliferation and neuronal and glial differentiation,” Dev. Neurobiol., vol. 75, no. 5, pp. 452–462, May 2015, doi: 10.1002/dneu.22235.

[6] D. A. Lim et al., “Chromatin remodelling factor Mll1 is essential for neurogenesis from postnatal neural stem cells,” Nature, vol. 458, no. 7237, pp. 529–533, Mar. 2009, doi: 10.1038/nature07726.

[7] J. M. Alarcón et al., “Chromatin acetylation, memory, and LTP are impaired in CBP+/-mice: a model for the cognitive deficit in Rubinstein-Taybi syndrome and its amelioration,” Neuron, vol. 42, no. 6, pp. 947–959, Jun. 2004, doi: 10.1016/j.neuron.2004.05.021.

[8] J. Guy, J. Gan, J. Selfridge, S. Cobb, and A. Bird, “Reversal of neurological defects in a mouse model of Rett syndrome,” Science, vol. 315, no. 5815, pp. 1143–1147, Feb. 2007, doi: 10.1126/science.1138389.

[9] H. T. Bjornsson et al., “Histone deacetylase inhibition rescues structural and functional brain deficits in a mouse model of Kabuki syndrome,” Sci. Transl. Med., vol. 6, no. 256, p. 256ra135, Oct. 2014, doi: 10.1126/scitranslmed.3009278.

[10] L. Zhang et al., “Inhibition of KDM1A activity restores adult neurogenesis and improves hippocampal memory in a mouse model of Kabuki syndrome,” Mol. Ther. Methods Clin. Dev., vol. 20, pp. 779–791, Mar. 2021, doi: 10.1016/j.omtm.2021.02.011.

[11] B. D. Yu, J. L. Hess, S. E. Horning, G. A. Brown, and S. J. Korsmeyer, “Altered Hox expression and segmental identity in Mll-mutant mice,” Nature, vol. 378, no. 6556, pp. 505–508, Nov. 1995, doi: 10.1038/378505a0.

[12] C. Kerimoglu et al., “KMT2A and KMT2B Mediate Memory Function by Affecting Distinct Genomic Regions,” Cell Rep., vol. 20, no. 3, pp. 538–548, Jul. 2017, doi: 10.1016/j.celrep.2017.06.072.

[13] C. N. Vallianatos et al., “Mutually suppressive roles of KMT2A and KDM5C in behaviour, neuronal structure, and histone H3K4 methylation,” *Commun*. Biol., vol. 3, no. 1, p. 278, Jun. 2020, doi: 10.1038/s42003-020-1001-6.

[14] J. A. Fahrner and H. T. Bjornsson, “Mendelian disorders of the epigenetic machinery: tipping the balance of chromatin states,” Annu. Rev. Genomics Hum. Genet., vol. 15, pp. 269–293, 2014, doi: 10.1146/annurev-genom-090613-094245.

[15] R. Terranova, H. Agherbi, A. Boned, S. Meresse, and M. Djabali, “Histone and DNA methylation defects at Hox genes in mice expressing a SET domain-truncated form of Mll,” Proc. Natl. Acad. Sci. U. S. A., vol. 103, no. 17, pp. 6629–6634, Apr. 2006, doi: 10.1073/pnas.0507425103.

[16] T. Reynisdottir, K. J. Anderson, L. Boukas, and H. T. Bjornsson, “Missense variants causing Wiedemann-Steiner syndrome preferentially occur in the KMT2A-CXXC domain and are accurately classified using AlphaFold2,” PLOS Genet., vol. 18, no. 6, p. e1010278, Jun. 2022, doi: 10.1371/journal.pgen.1010278.

[17] S. E. Sheppard et al., “Expanding the genotypic and phenotypic spectrum in a diverse cohort of 104 individuals with Wiedemann-Steiner syndrome,” Am. J. Med. Genet. A., vol. 185, no. 6, pp. 1649–1665, Jun. 2021, doi: 10.1002/ajmg.a.62124.

[18] E. Zemheri, P. Engin Zerk, and C. Ulukaya Durakbasa, “Calretinin immunohistochemical staining in Hirschsprung’s disease: An institutional experience,” North. Clin. Istanb., vol. 8, no. 6, pp. 601–606, 2021, doi: 10.14744/nci.2020.69376.

[19] K. B. Shpargel, J. Starmer, C. Wang, K. Ge, and T. Magnuson, “UTX-guided neural crest function underlies craniofacial features of Kabuki syndrome,” Proc. Natl. Acad. Sci., vol. 114, no. 43, pp. E9046–E9055, Oct. 2017, doi: 10.1073/pnas.1705011114.

[20] J. Schwenty-Lara, D. Nehl, and A. Borchers, “The histone methyltransferase KMT2D, mutated in Kabuki syndrome patients, is required for neural crest cell formation and migration,” Hum. Mol. Genet., vol. 29, no. 2, pp. 305–319, Jan. 2020, doi: 10.1093/hmg/ddz284.

[21] K. B. Shpargel, C. L. Mangini, G. Xie, K. Ge, and T. Magnuson, “The KMT2D Kabuki syndrome histone methylase controls neural crest cell differentiation and facial morphology,” Dev. Camb. Engl., vol. 147, no. 21, p. dev187997, Jul. 2020, doi: 10.1242/dev.187997.

[22] L. L. Baxter, L. Hou, S. K. Loftus, and W. J. Pavan, “Spotlight on spotted mice: a review of white spotting mouse mutants and associated human pigmentation disorders,” Pigment Cell Res., vol. 17, no. 3, pp. 215–224, Jun. 2004, doi: 10.1111/j.1600-0749.2004.00147.x.

[23] Z. Sarnyai, E. L. Sibille, C. Pavlides, R. J. Fenster, B. S. McEwen, and M. Toth, “Impaired hippocampal-dependent learning and functional abnormalities in the hippocampus in mice lacking serotonin(1A) receptors,” Proc. Natl. Acad. Sci. U. S. A., vol. 97, no. 26, pp. 14731–14736, Dec. 2000, doi: 10.1073/pnas.97.26.14731.

[24] A. K. Swonger and R. H. Rech, “Serotonergic and cholinergic involvement in habituation of activity and spontaneous alternation of rats in a Y maze,” J. Comp. Physiol. Psychol., vol. 81, no. 3, pp. 509–522, Dec. 1972, doi: 10.1037/h0033690.

[25] J. Boisgontier et al., “Anatomical and functional abnormalities on MRI in kabuki syndrome,” NeuroImage Clin., vol. 21, p. 101610, 2019, doi: 10.1016/j.nicl.2018.11.020.

[26] L. Li et al., “SoxD genes are required for adult neural stem cell activation,” Cell Rep., vol. 38, no. 5, p. 110313, Feb. 2022, doi: 10.1016/j.celrep.2022.110313.

[27] M. Durens, M. Soliman, J. Millonig, and E. DiCicco-Bloom, “Engrailed-2 is a cell autonomous regulator of neurogenesis in cultured hippocampal neural stem cells,” Dev. Neurobiol., vol. 81, no. 5, pp. 724–735, Jul. 2021, doi: 10.1002/dneu.22824.

[28] I. Schanze, D. Schanze, C. A. Bacino, S. Douzgou, B. Kerr, and M. Zenker, “Haploinsufficiency of SOX5, a member of the SOX (SRY-related HMG-box) family of transcription factors is a cause of intellectual disability,” Eur. J. Med. Genet., vol. 56, no. 2, pp. 108–113, Feb. 2013, doi: 10.1016/j.ejmg.2012.11.001.

[29] N. Gharani, R. Benayed, V. Mancuso, L. M. Brzustowicz, and J. H. Millonig, “Association of the homeobox transcription factor, ENGRAILED 2, 3, with autism spectrum disorder,” Mol. Psychiatry, vol. 9, no. 5, pp. 474–484, May 2004, doi: 10.1038/sj.mp.4001498.

[30] R. Ng, A. Kalinousky, J. A. Fahrner, H. T. Bjornsson, and J. Harris, “The social phenotype associated with Wiedemann-Steiner-syndrome: Autistic traits juxtaposed with high social drive and prosociality,” Am. J. Med. Genet. A., vol. 191, no. 10, pp. 2591–2601, Oct. 2023, doi: 10.1002/ajmg.a.63351.

[31] S. Banerjee-Basu and A. Packer, “SFARI Gene: an evolving database for the autism research community,” Dis. Model. Mech., vol. 3, no. 3–4, pp. 133–135, Mar. 2010, doi: 10.1242/dmm.005439.

[32] T. Gao, B. He, S. Liu, H. Zhu, K. Tan, and J. Qian, “EnhancerAtlas: a resource for enhancer annotation and analysis in 105 human cell/tissue types,” Bioinforma. Oxf. Engl., vol. 32, no. 23, pp. 3543–3551, Dec. 2016, doi: 10.1093/bioinformatics/btw495.

[33] N. T. Strande and J. S. Berg, “Defining the Clinical Value of a Genomic Diagnosis in the Era of Next-Generation Sequencing,” Annu. Rev. Genomics Hum. Genet., vol. 17, pp. 303–332, Aug. 2016, doi: 10.1146/annurev-genom-083115-022348.

[34] F. Morshedzadeh et al., “An Update on the Application of CRISPR Technology in Clinical Practice,” Mol. Biotechnol., pp. 1–19, Jun. 2023, doi: 10.1007/s12033-023-00724-z.

[35] J. Scharner and I. Aznarez, “Clinical Applications of Single-Stranded Oligonucleotides: Current Landscape of Approved and In-Development Therapeutics,” Mol. Ther., vol. 29, no. 2, pp. 540–554, Feb. 2021, doi: 10.1016/j.ymthe.2020.12.022.

[36] H. Liang, S. Hippenmeyer, and H. T. Ghashghaei, “A Nestin-cre transgenic mouse is insufficient for recombination in early embryonic neural progenitors,” Biol. Open, vol. 1, no. 12, pp. 1200–1203, Sep. 2012, doi: 10.1242/bio.20122287.

[37] S. A. Giusti et al., “Behavioral phenotyping of Nestin-Cre mice: implications for genetic mouse models of psychiatric disorders,” J. Psychiatr. Res., vol. 55, pp. 87–95, Aug. 2014, doi: 10.1016/j.jpsychires.2014.04.002.

[38] Y. L. Jung, C. Hung, J. Choi, E. A. Lee, and O. Bodamer, “Characterizing the molecular impact of KMT2D variants on the epigenetic and transcriptional landscapes in Kabuki syndrome,” Hum. Mol. Genet., vol. 32, no. 13, pp. 2251–2261, Jun. 2023, doi: 10.1093/hmg/ddad059.

[39] T. R. Luperchio et al., “Leveraging the Mendelian disorders of the epigenetic machinery to systematically map functional epigenetic variation,” eLife, vol. 10, p. e65884, Aug. 2021, doi: 10.7554/eLife.65884.

[40] G. Iacono et al., “Increased H3K9 methylation and impaired expression of Protocadherins are associated with the cognitive dysfunctions of the Kleefstra syndrome,” Nucleic Acids Res., vol. 46, no. 10, pp. 4950–4965, Jun. 2018, doi: 10.1093/nar/gky196.

[41] G. A. Carosso et al., “Precocious neuronal differentiation and disrupted oxygen responses in Kabuki syndrome,” JCI Insight, vol. 4, no. 20, pp. e129375, 129375, Oct. 2019, doi: 10.1172/jci.insight.129375.

[42] J. A. Fahrner et al., “Precocious chondrocyte differentiation disrupts skeletal growth in Kabuki syndrome mice,” JCI Insight, vol. 4, no. 20, pp. e129380, 129380, Oct. 2019, doi: 10.1172/jci.insight.129380.

[43] V. Faundes, G. Malone, W. G. Newman, and S. Banka, “A comparative analysis of KMT2D missense variants in Kabuki syndrome, cancers and the general population,” J. Hum. Genet., vol. 64, no. 2, pp. 161–170, Feb. 2019, doi: 10.1038/s10038-018-0536-6.

[44] H. T. Bjornsson et al., “Histone deacetylase inhibition rescues structural and functional brain deficits in a mouse model of Kabuki syndrome,” Sci. Transl. Med., vol. 6, no. 256, p. 256ra135, Oct. 2014, doi: 10.1126/scitranslmed.3009278.

[45] C. Galichet, R. Lovell-Badge, and K. Rizzoti, “Nestin-Cre mice are affected by hypopituitarism, which is not due to significant activity of the transgene in the pituitary gland,” PloS One, vol. 5, no. 7, p. e11443, Jul. 2010, doi: 10.1371/journal.pone.0011443.

[46] B. Paigen, A. Morrow, C. Brandon, D. Mitchell, and P. Holmes, “Variation in susceptibility to atherosclerosis among inbred strains of mice,” Atherosclerosis, vol. 57, no. 1, pp. 65–73, Oct. 1985, doi: 10.1016/0021-9150(85)90138-8.

[47] S. Hayashi, P. Lewis, L. Pevny, and A. P. McMahon, “Efficient gene modulation in mouse epiblast using a Sox2Cre transgenic mouse strain,” Mech. Dev., vol. 119 Suppl 1, pp. S97–S101, Dec. 2002, doi: 10.1016/s0925-4773(03)00099-6.

[48] F. Tronche et al., “Disruption of the glucocorticoid receptor gene in the nervous system results in reduced anxiety,” Nat. Genet., vol. 23, no. 1, pp. 99–103, Sep. 1999, doi: 10.1038/12703.

[49] A. Ventura et al., “Restoration of p53 function leads to tumour regression in vivo,” Nature, vol. 445, no. 7128, pp. 661–665, Feb. 2007, doi: 10.1038/nature05541.

[50] D. N. Feather-Schussler and T. S. Ferguson, “A Battery of Motor Tests in a Neonatal Mouse Model of Cerebral Palsy,” J. Vis. Exp. JoVE, no. 117, p. 53569, Nov. 2016, doi: 10.3791/53569.

[51] A.-K. Kraeuter, P. C. Guest, and Z. Sarnyai, “The Y-Maze for Assessment of Spatial Working and Reference Memory in Mice,” Methods Mol. Biol. Clifton NJ, vol. 1916, pp. 105–111, 2019, doi: 10.1007/978-1-4939-8994-2_10.

[52] A. Fedorov et al., “3D Slicer as an image computing platform for the Quantitative Imaging Network,” Magn. Reson. Imaging, vol. 30, no. 9, pp. 1323–1341, Nov. 2012, doi: 10.1016/j.mri.2012.05.001.

[53] Adams D, Collyer M, Kaliontzopoulou A, Baken E, “Geomorph: Software for geometric morphometric analyses. R package version 4.0.7.” 2024. [Online]. Available: https://cran.r-project.org/package=geomorph.

[54] C. P. Klingenberg, M. Barluenga, and A. Meyer, “Shape Analysis of Symmetric Structures: Quantifying Variation Among Individuals and Asymmetry,” Evolution, vol. 56, no. 10, pp. 1909–1920, 2002, doi: 10.1111/j.0014-3820.2002.tb00117.x.

[55] P. Gunz, P. Mitteroecker, S. Neubauer, G. W. Weber, and F. L. Bookstein, “Principles for the virtual reconstruction of hominin crania,” J. Hum. Evol., vol. 57, no. 1, pp. 48–62, Jul. 2009, doi: 10.1016/j.jhevol.2009.04.004.

[56] J. Schindelin et al., “Fiji: an open-source platform for biological-image analysis,” Nat. Methods, vol. 9, no. 7, pp. 676–682, Jul. 2012, doi: 10.1038/nmeth.2019.

[57] S. N. Bernas, O. Leiter, T. L. Walker, and G. Kempermann, “Isolation, Culture and Differentiation of Adult Hippocampal Precursor Cells,” Bio-Protoc., vol. 7, no. 21, p. e2603, Nov. 2017, doi: 10.21769/BioProtoc.2603.

[58] N. L. Bray, H. Pimentel, P. Melsted, and L. Pachter, “Near-optimal probabilistic RNA-seq quantification,” Nat. Biotechnol., vol. 34, no. 5, pp. 525–527, May 2016, doi: 10.1038/nbt.3519.

[59] F. Ramírez et al., “deepTools2: a next generation web server for deep-sequencing data analysis,” Nucleic Acids Res., vol. 44, no. W1, pp. W160–165, Jul. 2016, doi: 10.1093/nar/gkw257.

[60] W. J. Kent et al., “The human genome browser at UCSC,” Genome Res., vol. 12, no. 6, pp. 996–1006, Jun. 2002, doi: 10.1101/gr.229102.

[61] R Core Team, “R: A language and environment for statistical computing.” R Foundation for Statistical Computing, Vienna, Austria., 2021. [Online]. Available: https://www.R-project.org/

[62] C. Soneson, M. I. Love, and M. D. Robinson, “Differential analyses for RNA-seq: transcript-level estimates improve gene-level inferences,” F1000Research, vol. 4, p. 1521, 2015, doi: 10.12688/f1000research.7563.2.

[63] S. Durinck, P. T. Spellman, E. Birney, and W. Huber, “Mapping identifiers for the integration of genomic datasets with the R/Bioconductor package biomaRt,” Nat. Protoc., vol. 4, no. 8, pp. 1184–1191, 2009, doi: 10.1038/nprot.2009.97.

[64] M. I. Love, W. Huber, and S. Anders, “Moderated estimation of fold change and dispersion for RNA-seq data with DESeq2,” Genome Biol., vol. 15, no. 12, p. 550, 2014, doi: 10.1186/s13059-014-0550-8.

[65] K. Strimmer, “fdrtool: a versatile R package for estimating local and tail area-based false discovery rates,” Bioinforma. Oxf. Engl., vol. 24, no. 12, pp. 1461–1462, Jun. 2008, doi: 10.1093/bioinformatics/btn209.

[66] Y. Liao, J. Wang, E. J. Jaehnig, Z. Shi, and B. Zhang, “WebGestalt 2019: gene set analysis toolkit with revamped UIs and APIs,” Nucleic Acids Res., vol. 47, no. W1, pp. W199–W205, Jul. 2019, doi: 10.1093/nar/gkz401.

[67] Z. Gu, R. Eils, and M. Schlesner, “Complex heatmaps reveal patterns and correlations in multidimensional genomic data,” Bioinforma. Oxf. Engl., vol. 32, no. 18, pp. 2847– 2849, Sep. 2016, doi: 10.1093/bioinformatics/btw313.

[68] “The Landscape of Isoform Switches in Human Cancers-PubMed.” Accessed: May 13, 2024. [Online]. Available: https://pubmed.ncbi.nlm.nih.gov/28584021/

[69] T. Hulsen, “DeepVenn – a web application for the creation of area-proportional Venn diagrams using the deep learning framework Tensorflow.js.” Department of Hospital Services & Informatics, Philips Research, Eindhoven, the Netherlands, 2022.

[70] Shen L, Sinai ISoMaM, “GeneOverlap: Test and visualize gene overlaps.” 2024.

[71] B. Langmead and S. L. Salzberg, “Fast gapped-read alignment with Bowtie 2,” Nat. Methods, vol. 9, no. 4, pp. 357–359, Mar. 2012, doi: 10.1038/nmeth.1923.

[72] H. Li et al., “The Sequence Alignment/Map format and SAMtools,” Bioinforma. Oxf. Engl., vol. 25, no. 16, pp. 2078–2079, Aug. 2009, doi: 10.1093/bioinformatics/btp352.

[73] Y. Zhang et al., “Model-based analysis of ChIP-Seq (MACS),” Genome Biol., vol. 9, no. 9, p. R137, 2008, doi: 10.1186/gb-2008-9-9-r137.

[74] R. Stark and G. Brown, “DiffBind: Differential binding analysis of ChIP-Seq peak data”.

[75] “Exploring Epigenomic Datasets by ChIPseeker-PubMed.” Accessed: May 13, 2024. [Online]. Available: https://pubmed.ncbi.nlm.nih.gov/36286622/

[76] H. Li, “Minimap2: pairwise alignment for nucleotide sequences,” Bioinforma. Oxf. Engl., vol. 34, no. 18, pp. 3094–3100, Sep. 2018, doi: 10.1093/bioinformatics/bty191.

[77] J. T. Robinson et al., “Integrative genomics viewer,” Nat. Biotechnol., vol. 29, no. 1, pp. 24–26, Jan. 2011, doi: 10.1038/nbt.1754.

