## Supplementary figures and images for "*In-utero* rescue of neurological dysfunction in a mouse model of Wiedemann-Steiner syndrome"

### Supplemental Figure 2

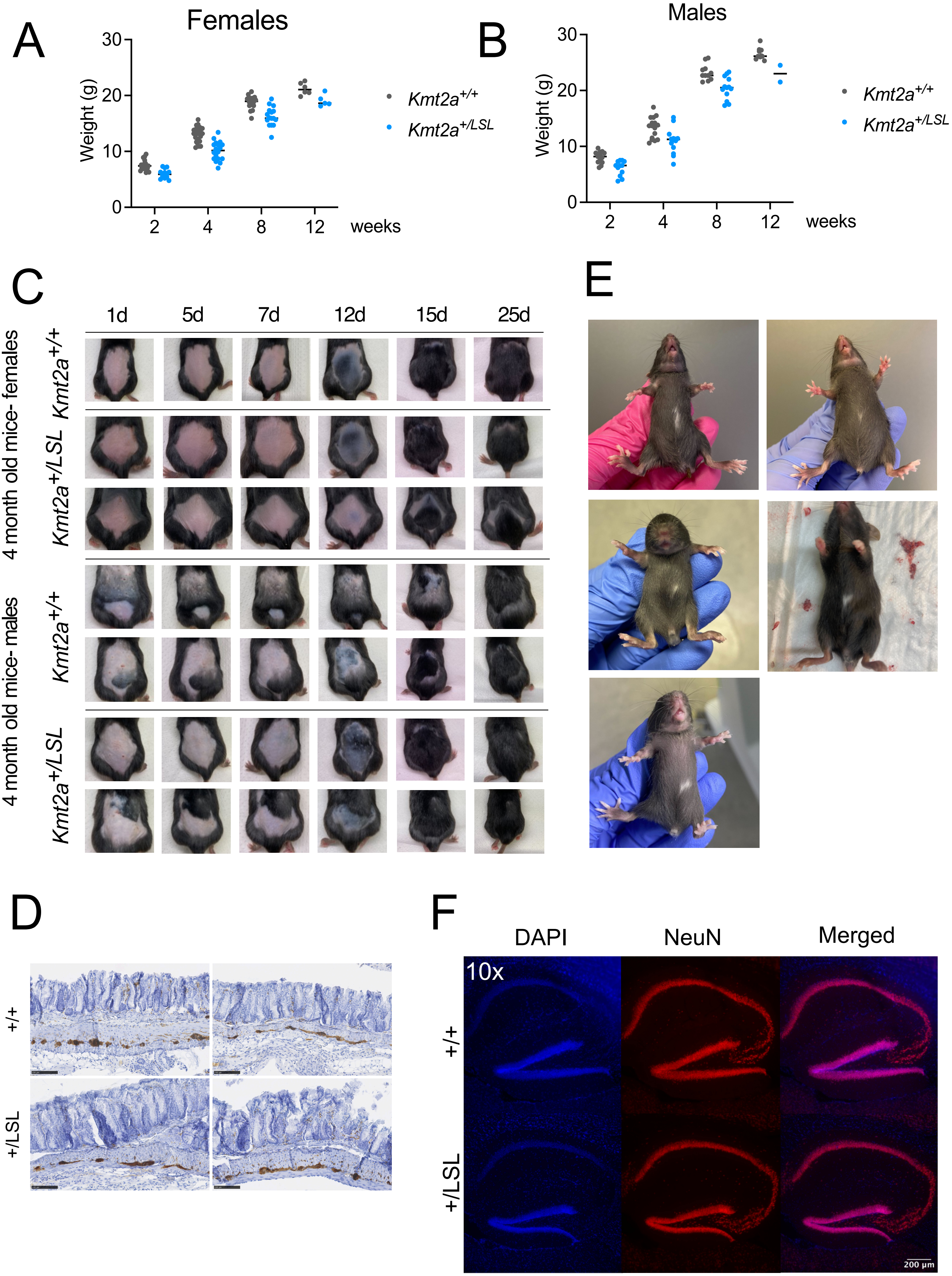

### Supplemental Figure 3

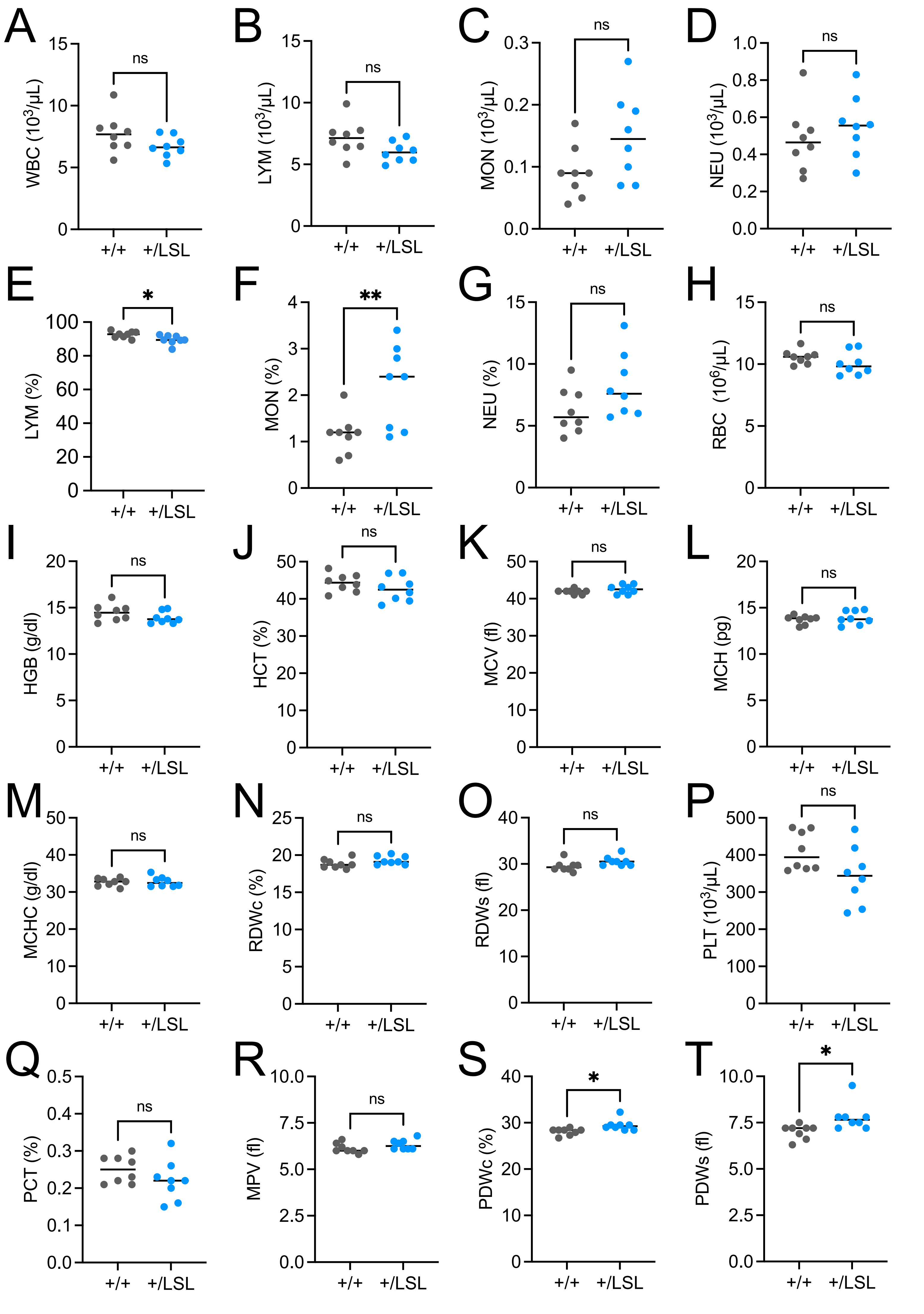

### Supplemental Figure 4

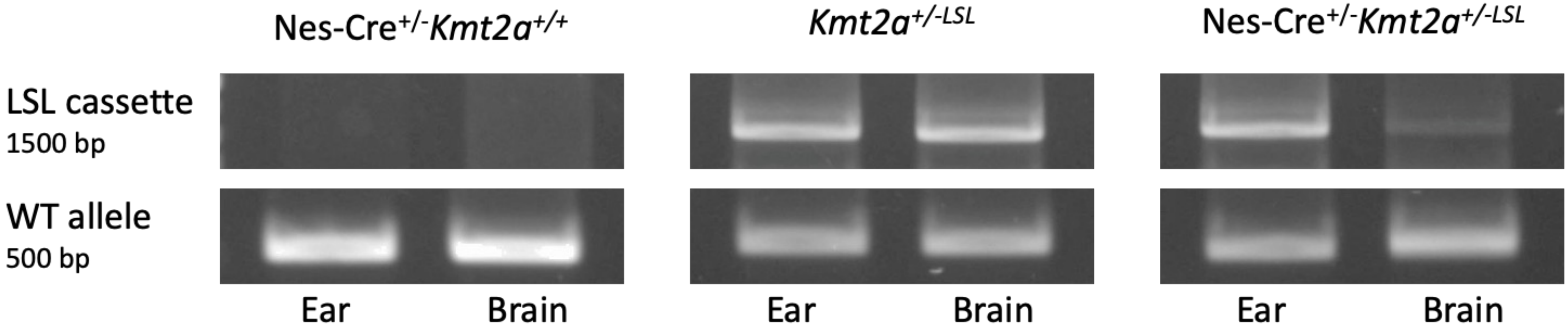

### Supplemental Figure 5

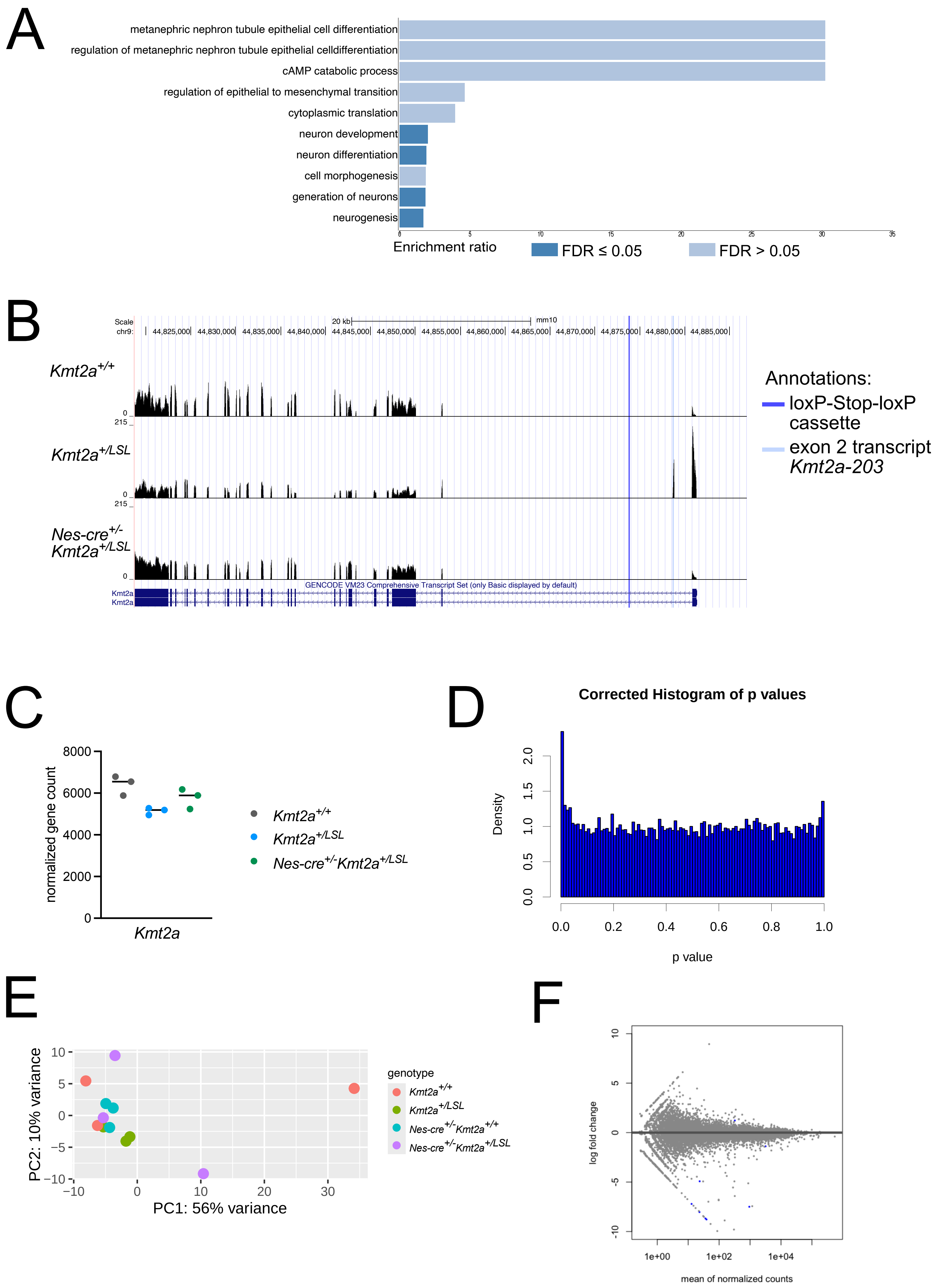

### Supplemental Figure 6

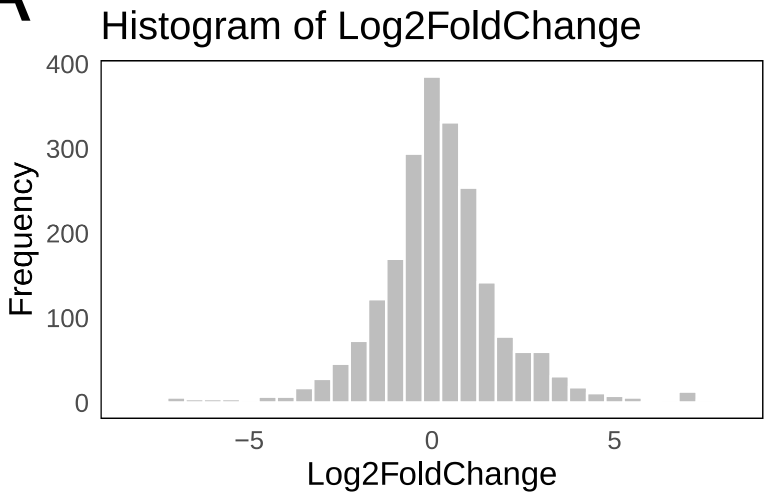

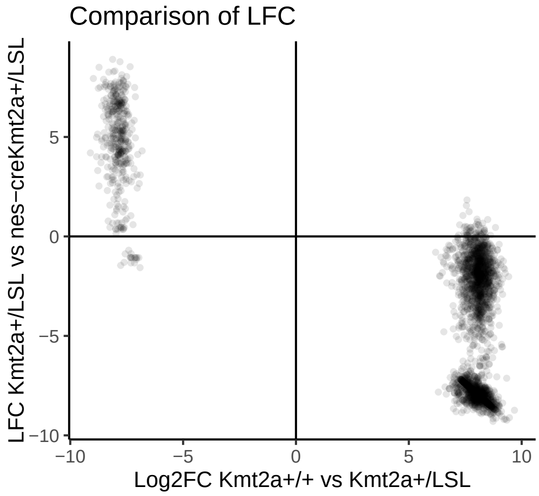

### Supplemental Figure 7

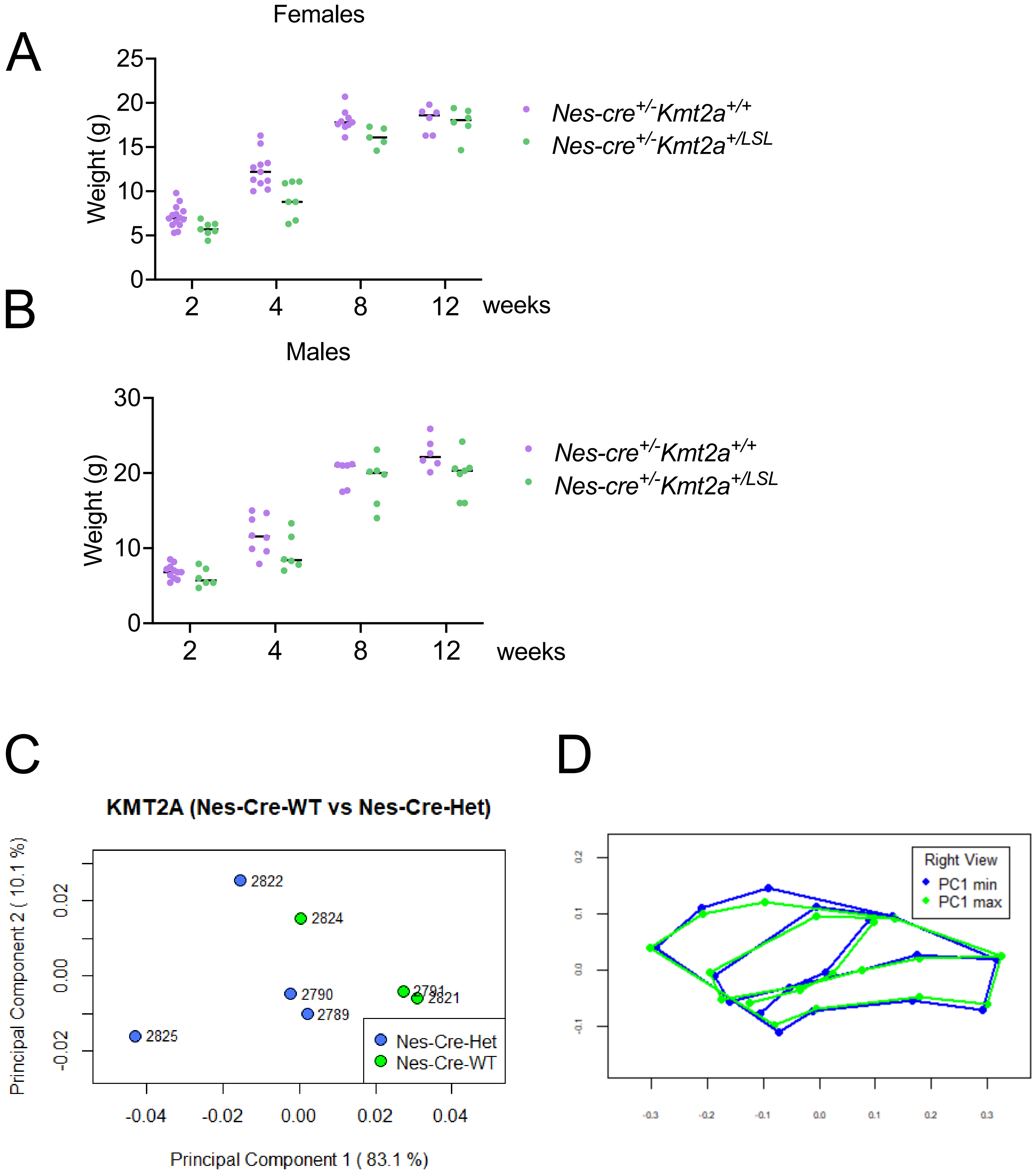
