## Supplemental Figure 1 for "*In-utero* rescue of neurological dysfunction in a mouse model of Wiedemann-Steiner syndrome"

A

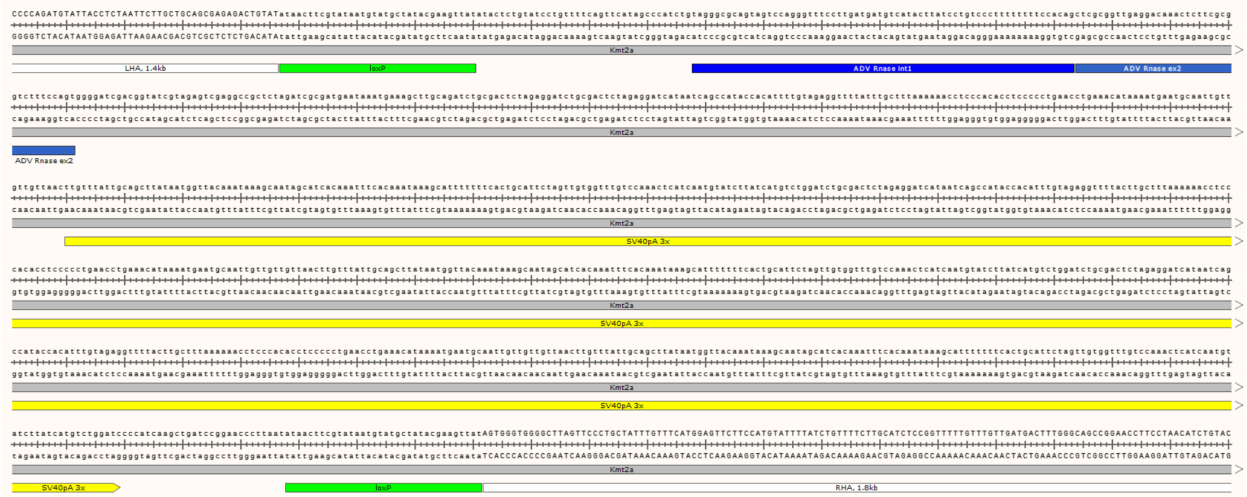


B

| crRNA | Sequence | PAM | Score | Locus | Gene |
|---|---|---|---|---|---|
| Kmt2a_in1_crRNA1: | CTGCAGCGAGAGACTGTATG | AGG | 100.0 | Chr9: - 44873762 | *Kmt2a* |
| CTGCAGCGAGAGACTGTATG | TTGGAGCCAGAGACTGTATG | AAG | 2.8 | Chr 3: +60581077 | *Mbnl1* |
| Kmt2a_in1_crRNA2: | GAGACTGTATGAGGTATCAG | TGG | 100.0 | Chr9: 44873753 | *Kmt2a* |
| GAGACTGTATGAGGTATCAG | AAGACTGTATGAAGTATCAG | AAG | 3.9 | Chr4: -59081019 | *Shoc1* |

C

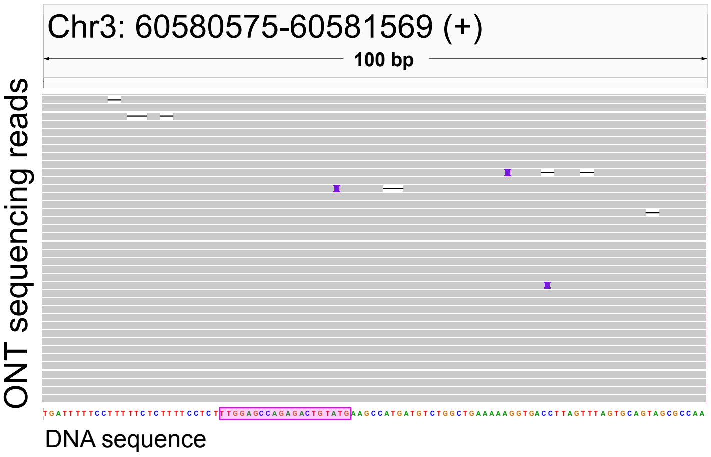


D

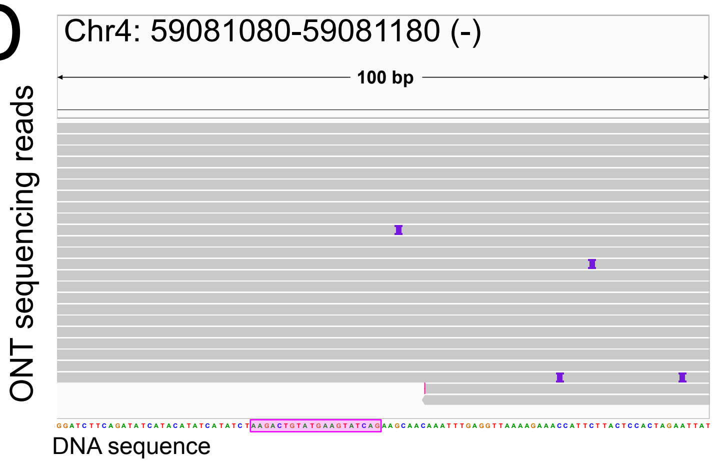


E

| | Het x Het | | | |
|---|---|---|---|---|
| 35724 | WT | Het | Hom | |
| Female | 8 | 20 | 0 | |
| Male | 14 | 12 | 0 | |
| Total | 22 | 32 | 0 | 54 |
| % | 40.7 | 59.3 | 0.0 | |

F

| | Het x Het | | | |
|---|---|---|---|---|
| 35725 | WT | Het | Hom | |
| Female | 11 | 22 | 0 | |
| Male | 13 | 13 | 0 | |
| Total | 24 | 35 | 0 | 59 |
| % | 40.7 | 59.3 | 0.0 | |

G

| | Het x Het | | | |
|---|---|---|---|---|
| 35727 | wt | het | Hom | |
| female | 6 | 10 | 2 | |
| male | 3 | 11 | 3 | |
| total | 9 | 21 | 5 | 35 |
| % | 25.7 | 60.0 | 14.3 | p, ns |
